# Aberrant lipid accumulation and retinal pigmental epithelium dysfunction in PRCD-deficient mice

**DOI:** 10.1101/2024.03.08.584131

**Authors:** Sree I. Motipally, Douglas R. Kolson, Tongju Guan, Saravanan Kolandaivelu

## Abstract

Progressive Rod-Cone Degeneration (PRCD) is an integral membrane protein found in photoreceptor outer segment (OS) disc membranes and its function remains unknown. Mutations in *Prcd* are implicated in *Retinitis pigmentosa* (RP) in humans and multiple dog breeds. PRCD-deficient models exhibit decreased levels of cholesterol in the plasma. However, potential changes in the retinal cholesterol remain unexplored. In addition, impaired phagocytosis observed in these animal models points to potential deficits in the retinal pigment epithelium (RPE). Here, using a *Prcd^-/-^*murine model we investigated the alterations in the retinal cholesterol levels and impairments in the structural and functional integrity of the RPE. Lipidomic and immunohistochemical analyses show a 5-fold increase in the levels of cholesteryl esters (C.Es) and accumulation of neutral lipids in the PRCD-deficient retina, respectively, indicating alterations in total retinal cholesterol. Longitudinal fundus and spectral domain optical coherence tomography (SD-OCT) examinations showed focal lesions and RPE hyperreflectivity. Strikingly, the RPE of *Prcd^-/-^* mice exhibited age-related pathological features such as neutral lipid deposits, lipofuscin accumulation, Bruch’s membrane (BrM) thickening and drusenoid focal deposits, mirroring an Age-related Macular Degeneration (AMD)-like phenotype. We propose that the extensive lipofuscin accumulation likely impairs lysosomal function, leading to the defective phagocytosis observed in *Prcd^-/-^*mice. Our findings support the dysregulation of retinal cholesterol homeostasis in the absence of PRCD. Further, we demonstrate that progressive photoreceptor degeneration in *Prcd^-/-^*mice is accompanied by progressive structural and functional deficits in the RPE, which likely exacerbates vision loss over time.

## Introduction

Progressive rod-cone degeneration (PRCD) is a highly conserved photoreceptor OS specific protein with an unknown function that strongly associates with the disc membrane (1). In humans, mutations in *Prcd* are implicated in *Retinitis pigmentosa* (RP) - a highly heterogeneous hereditary retinal dystrophy characterized by progressive degeneration of rod photoreceptors followed by cone loss (2). The most common mutation in *Prcd,* Cys2Tyr (C2Y), is associated with late-onset autosomal recessive photoreceptor degeneration (3). The C2Y mutation abrogates palmitoylation rendering the protein unstable and mis-localized to the inner segment (IS) (4). Both canine *Prcd^C2Y/C2Y^* and mouse *Prcd^-/-^* models exhibit disorganized OS structure, uneven disc diameters and vesicular profiles (5–7). Studies in canine *Prcd^C2Y/C2Y^* models also reported a ∼50% reduction in OS disc renewal rates (5,8). Recent study demonstrated defective flattening of nascent discs in *Prcd^-/-^* mice, suggesting a critical role for PRCD in photoreceptor OS disc morphogenesis (7). However, the precise role of PRCD in maintaining OS structure and function remains obscure.

A distinct composition of proteins and lipids are necessary for establishing proper OS disc structure and function (9,10). The membrane-rich rod OS is composed of equal proportions of proteins and lipids by weight (11). Phospholipids and cholesterol account for ∼95% (w/w) and ∼5% (w/w) of the total rod OS lipid content, respectively (12). Docosahexaenoic acid (DHA, 22:6n-3) accounts for ∼50% of the total phospholipid fatty acid in the rod OS membranes and is critical for the maintenance of rod OS membrane fluidity, renewal, and rod function. (11,13–15). Furthermore, phospholipid composition of the rod OS can regulate the cholesterol content of the disc membranes (16). Decreased levels of DHA in the rod OS were previously reported in canine *Prcd^C2Y/C2Y^* and various other models of RP (17,18). Although reduced levels of plasma cholesterol were reported earlier in canine *Prcd^C2Y/C2Y^* models (19), to date, no study has investigated the potential changes in retinal cholesterol in *Prcd-*mutant models.

Photoreceptor OS membranes, including DHA and cholesterol, are recycled as a part of the daily process of phagocytosis of photoreceptor distal ends by the RPE. (20–23). Although no changes were found in the rate of synthesis and release of DHA by the RPE in *Prcd^C2Y/C2Y^* dogs (24), a recent study shows impaired phagocytosis in *Prcd*^-/-^ mice (25). Therefore, we sought to investigate alterations in retinal cholesterol levels and phospholipid composition in *Prcd^-/-^* mice in the current study. Impaired phagocytosis, which is also observed in other models of inherited retinal degenerations (26,27), can lead to RPE dysfunction and death (28). The structural and functional integrity of the RPE in *Prcd*^-/-^ models remain understudied. To this end, we performed a comprehensive assessment of RPE health in *Prcd*^-/-^ mice.

We used global *Prcd^-/-^* mice to investigate both the alterations in the retinal cholesterol levels and characterize the potential pathophysiological changes in the RPE that can contribute to the observed photoreceptor deficits. We found elevated levels of C.Es and neutral lipid accumulation in the PRCD-deficient retina which indicate alterations in total cholesterol. Interestingly, *Prcd*^-/-^ mice exhibit thickened BrM, subretinal and sub-RPE drusenoid focal deposits mimicking an Age-related Macular Degeneration (AMD)-like phenotype. Furthermore, we found extensive accumulation of lipofuscin and propose that it likely leads to RPE lysosomal dysfunction, which may underlie the impaired phagocytosis observed in *Prcd*^-/-^ mice. In conclusion, we report multiple lines of evidence that support dysregulation of retinal cholesterol homeostasis and RPE impairment in *Prcd*^-/-^ mice. We suggest that the RPE deficits observed in *Prcd*^-/-^ mice may exacerbate photoreceptor dysfunction and degeneration. Our report is the first comprehensive study of the structural and functional anomalies that occur in the RPE of *Prcd*^-/-^ mice.

## Results

### PRCD-deficient retina accumulates cholesteryl esters

As the phospholipid composition can regulate the cholesterol content of the disc membranes (16), we hypothesized that the reduced levels of DHA in the PRCD-deficient retina (17) may lead to alterations in the total levels of retinal cholesterol. To investigate this possibility, we used untargeted LC-MS/MS to quantify the global retinal lipid composition in *Prcd^-/-^* mice and wildtype (WT) controls at 2-months postnatal age. We chose to quantify the global alterations in the retinal lipidome instead of exclusively the rod OS to avoid the potential loss of lipid content arising from the extracellular vesicles (EVs) that form in the absence of PRCD (7); ( Fig. S1). We would like to emphasize that no histological evidence of retinal cell death was observed in 2-month-old *Prcd^-/-^*mice that can account for the observed lipid alterations. The relative abundance of various phosphatidyl choline (PC) and phosphatidyl ethanolamine (PE) species were significantly lower in *Prcd^-/-^* retina when compared to WT controls (Fig. 1B, C). First, the levels of various DHA-PC and DHA-PE species were reduced in *Prcd^-/-^*retina (Fig. 1B, C; *yellow highlights*). This finding is consistent with previous findings that reported reduced DHA levels in the rod OS of *Prcd^C2Y/C2Y^* canines (17). *Prcd^-/-^* retina also showed decreased levels of lyso PC, a blood-derived source of retinal DHA, (Fig. 1B inset) and lyso PE species (Fig. 1C inset) between the groups. Thirdly, the overall abundance of long-chain sphingomyelin (SM) species was significantly higher in *Prcd^-/-^* retina when compared to WT controls (Fig. 1D). Lastly and most notably, *Prcd^-/-^* retinas showed a 5-fold increase in the levels of cholesteryl esters (C.Es) when compared to WT controls (Fig. 1E). However, no significant changes were observed in the levels of free cholesterol (FC) (Fig. S2A) and triacylglycerols (TAGs) (Fig. S2B) between the groups. Altogether, in addition to the alterations in total retinal cholesterol, these data support diminished levels of two major glycerophospholipids - PC and PE, and suggests sphingolipid dysregulation in PRCD-deficient retinas.

**Figure 1.**
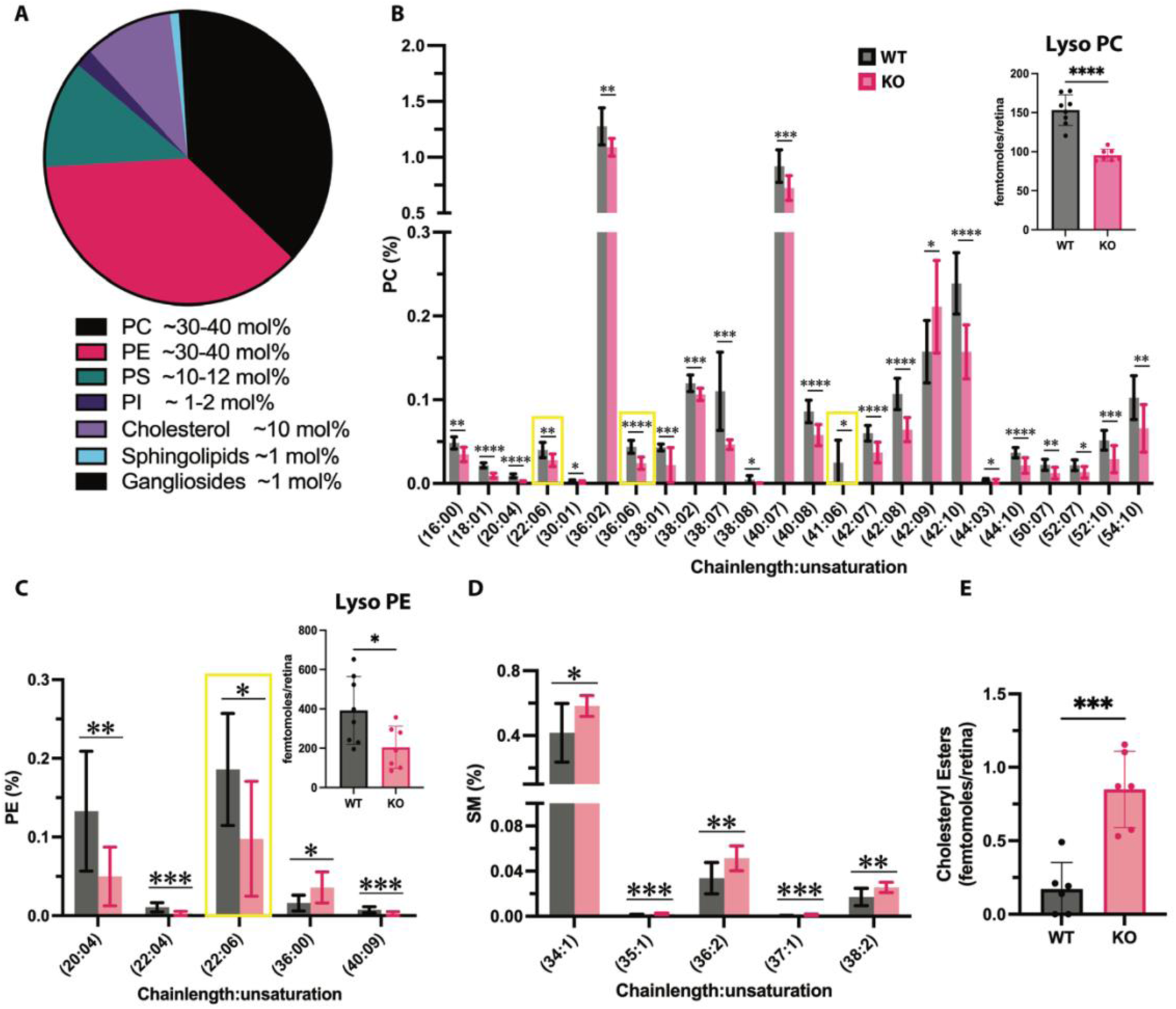
Lipid alterations in PRCD-deficient retina. Untargeted LC-MS/MS analysis of whole retinas from *Prcd^-/-^*and WT mice at P60 (WT, n=8; KO, n=7 mice). **A.** Pie chart illustrates the lipid composition of the rod photoreceptor OS in the bovine retina (10). **B.** The relative abundance of Phosphatidyl choline (PC) species between *Prcd^-/-^* and WT. The inset shows the total amount of lyso-PC species detected between the groups. **C.** The relative abundance of Phosphatidyl ethanolamine (PE) species. The inset shows the total amount of lyso-PE species detected. **D.** Relative abundance of sphingomyelin (SM) species between the groups. All lipid subspecies are shown as a percentage of total lipid classes detected. **E.** Quantification of C.Es between the groups (n=6). Groups were compared using either unpaired t-tests or unpaired multiple t-tests. No corrections for multiple comparisons were made. Data are represented as mean ± SD. Significance values are indicated as follows - *P<0.05; **P<0.02; ***P<0.001; ****P<0.0001.

### *Prcd^-/-^* mice exhibit neutral lipid accumulation in the photoreceptor IS and OS

To further investigate altered cholesterol levels observed in PRCD-deficient retinas, we asked whether C.Es are accumulating within the photoreceptor cells since PRCD is exclusively expressed in the photoreceptors (1,4). To test this possibility, we used Nile Red, a widely used fluorescent dye to detect neutral lipids that include FC and C.Es (29), on the retinal cryosections obtained from *Prcd^-/-^* and WT mice, at 2- and 4-month postnatal ages. While no differences were observed in the staining at 2-months, we found accumulation of lipid deposits in the photoreceptor IS and OS (Fig. 2*, B, B’*, *arrows and arrowheads*) of 4-month-old *Prcd^-/-^* retinas that were absent in age-matched WT controls (Fig. 2, *A*, *A’*). We next hypothesized that excess C.Es in *Prcd^-/-^* mice may lead to lipid droplet (LD) accumulation as LDs are universal storage organelles for neutral lipids (30). LDs are composed of a monolayer of amphipathic lipids and proteins encompassing a neutral lipid core that primarily consists of C.Es and TAGs (31,32). To check this, we investigated the expression of adipocyte differentiation-related protein/perilipin 2 (ADRP/PLIN2), which is exclusively localized to the surface of LDs (30). As expected, *Prcd^-/-^* mice showed significantly higher ADRP/PLIN2 expression in the photoreceptor IS (Fig. 2D, *D’, E*) and the OS (Fig. 2D, D’, F) when compared to age-matched WT controls (Fig. 2C, C’, E, F). These data indicate that, in addition to EVs, *Prcd^-/-^* retina may also show an accumulation of LDs, which further corroborates altered retinal cholesterol levels in the PRCD-deficient retina.

**Figure 2.**
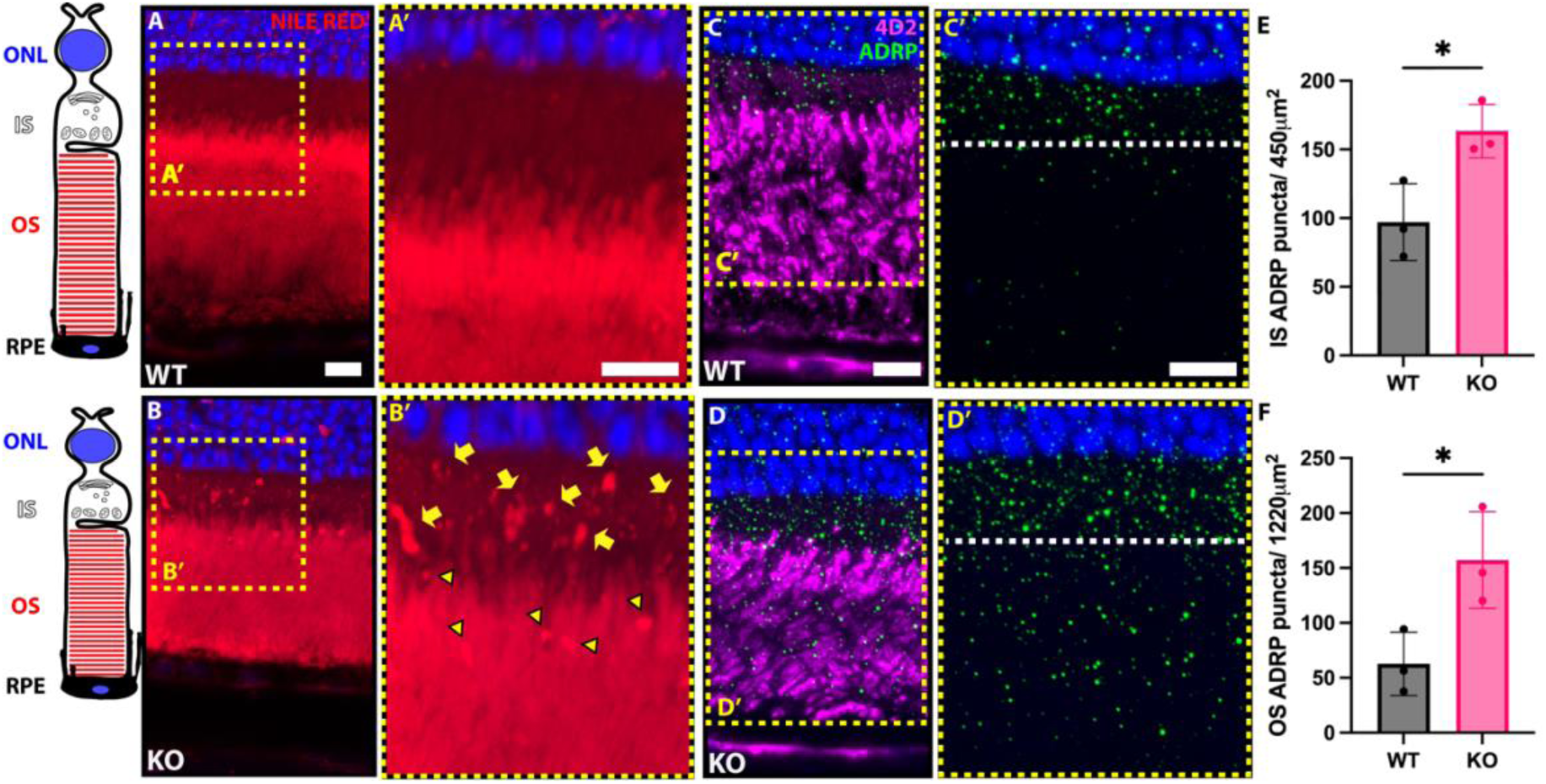
Accumulation of neutral lipids and ADRP overexpression in the IS and OS of PRCD-deficient photoreceptors. **A-B**. Neutral lipid staining (*red*) of retinal cross-sections from 4-month-old *Prcd^-/-^* and WT mice using Nile Red. Nuclei are stained with DAPI (*blue*). The photoreceptor schematic on the left is aligned to reflect the location of various photoreceptor compartments. Insets **A’** and **B’** are magnified to show neutral lipid accumulation. Yellow arrows and arrowheads in B’ point to lipid deposits in the IS and OS, respectively, in *Prcd^-/-^* mice. **C-D**. Retinal cross-sections from 4-month-old *Prcd^-/-^*and WT mice stained with lipid droplet specific protein – ADRP/PLIN2 (*green*). Insets **C’** and **D’** show the magnified view of ADRP puncta. The white dashed line indicates the IS-OS junction. **E, F.** Quantification of ADRP puncta in the IS and OS from control and *Prcd^-/-^*mice (n=3 biological replicates, 3 technical replicates; *P<0.05). Groups were compared using two-tailed Student t-test. Data are represented as mean ± SD. Scale bars -10µm. RPE – Retinal pigmental epithelium, OS – Outer segment, IS – Inner segment, ONL-outer nuclei layer.

### *Prcd*^-/-^ mice exhibit AMD-like pathological features

Next, we performed fundoscopy to investigate and track potential RPE abnormalities in *Prcd^-/-^* mice along with age-matched WT controls beginning from 2 to 10 months of age at 1-month intervals. Strikingly, we observed the presence of yellow-white punctate lesions with sharp borders as early as ∼2 months in *Prcd^-/-^*mice, and their numbers and size progressively increased with age (Fig. 3B, D, F). These lesions were absent in WT control mice across all the tested ages (Fig. 3A, C, E). Further evaluation by spectral-domain optical coherence tomography (SD-OCT) revealed substantial hyperreflective foci (HRF), that progressively increased in number in *Prcd^-/-^* mice (Fig. 3J; arrows) when compared to WT controls (Fig. 3I; *arrows*). Interestingly, we observed a loss of laminar delineation between the ellipsoid zone (EZ) and interdigitation zone (IZ) bands (Fig. 3H, *square brackets*) that is represented by an overall increased reflectivity in the photoreceptor OS region in *Prcd^-/-^* mice (Fig. 3K) when compared to WT controls (Fig. 3G, K).

**Figure 3.**
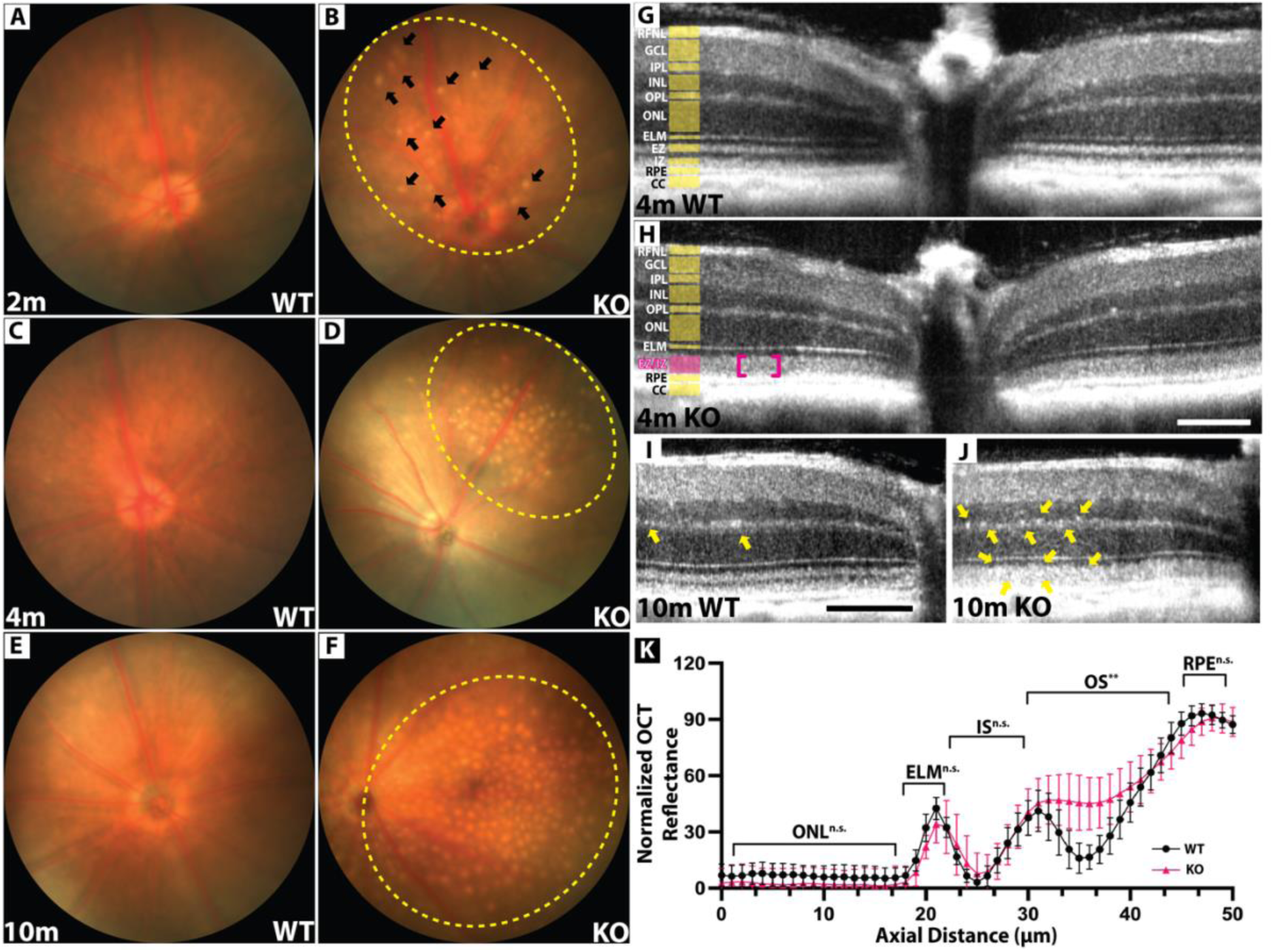
*Prcd*^-/-^ mice show yellowish punctate lesions and altered EZ-IZ laminar delineation. **A-F.** Fundus images of control (A, C, E) and *Prcd^-/-^* mice (B, D, F) at 2- (n=10),4- (n=7), and 10- (n=3) months postnatal ages showing yellow-white lesions (*black arrows/ dashed yellow ovals*). **G-I.** In vivo SD-OCT structural evaluation of WT control (G) and *Prcd^-/-^* mice (H) at 4 months (n=11 per group). The relative thickness of various OCT bands is demarcated by yellow highlights, and the loss of laminar delineation between EZ and IZ bands in *Prcd^-/-^* mice is indicated by magenta brackets (H)**. I-J.** Representative SD-OCT images from 10-month-old WT and *Prcd^-/-^* mice showing the overall reflectivity of various bands. *Prcd^-/-^*mice show hyperreflectivity in EZ/IZ bands and hyperreflective foci (J, *yellow arrows*) in the EZ/IZ and OPL bands (n=3). **K.** Averaged and normalized OCT mean reflectance signal from middle/outer retina vs axial tissue depth in 4-month-old mice. Groups were compared with multiple unpaired t-tests with Welch correction (n=5, P-value range for OS=0.01-0.03). RFNL – retinal nerve fiber layer; GCL-ganglion cell layer; IPL-inner plexiform layer; INL– inner nuclear layer; OPL – outer plexiform layer; ONL-outer nuclei layer; ELM-external limiting membrane; EZ – ellipsoid zone; IZ – interdigitating zone; RPE-retinal pigment epithelium; CC-choriocapillaris. Scale bars -100µm.

To characterize the punctate lesions observed in the fundus, we performed Hematoxylin and Eosin (H&E) histological staining on serial plastic sections obtained from 4-month-old *Prcd^-/-^*and WT control mice. In contrast to WT controls (Fig. 4C), we observed the presence of focal deposits resembling drusen (Fig. 4A and Fig. S3*A*, *black arrows*) and subretinal drusenoid deposits (SDD) (Fig. 4B, *dashed ovals*; Fig. S3*B*, *white arrows*) in PRCD-deficient retinas. In addition, we observed RPE thinning and atrophy in patches (Fig. 4D, E, *arrows*), RPE sloughing (Fig. 4D, E, *arrowheads*), RPE-free zones (Fig. 4D, E, *stars*), as well as the presence of non-pigmented cells in the subretinal space (Fig. 4F, *arrows*) in *Prcd^-/-^* retinas. Furthermore, these subretinal cells showed a high expression of apolipoprotein E (ApoE) (Fig. S3*C, D*).

**Figure 4.**
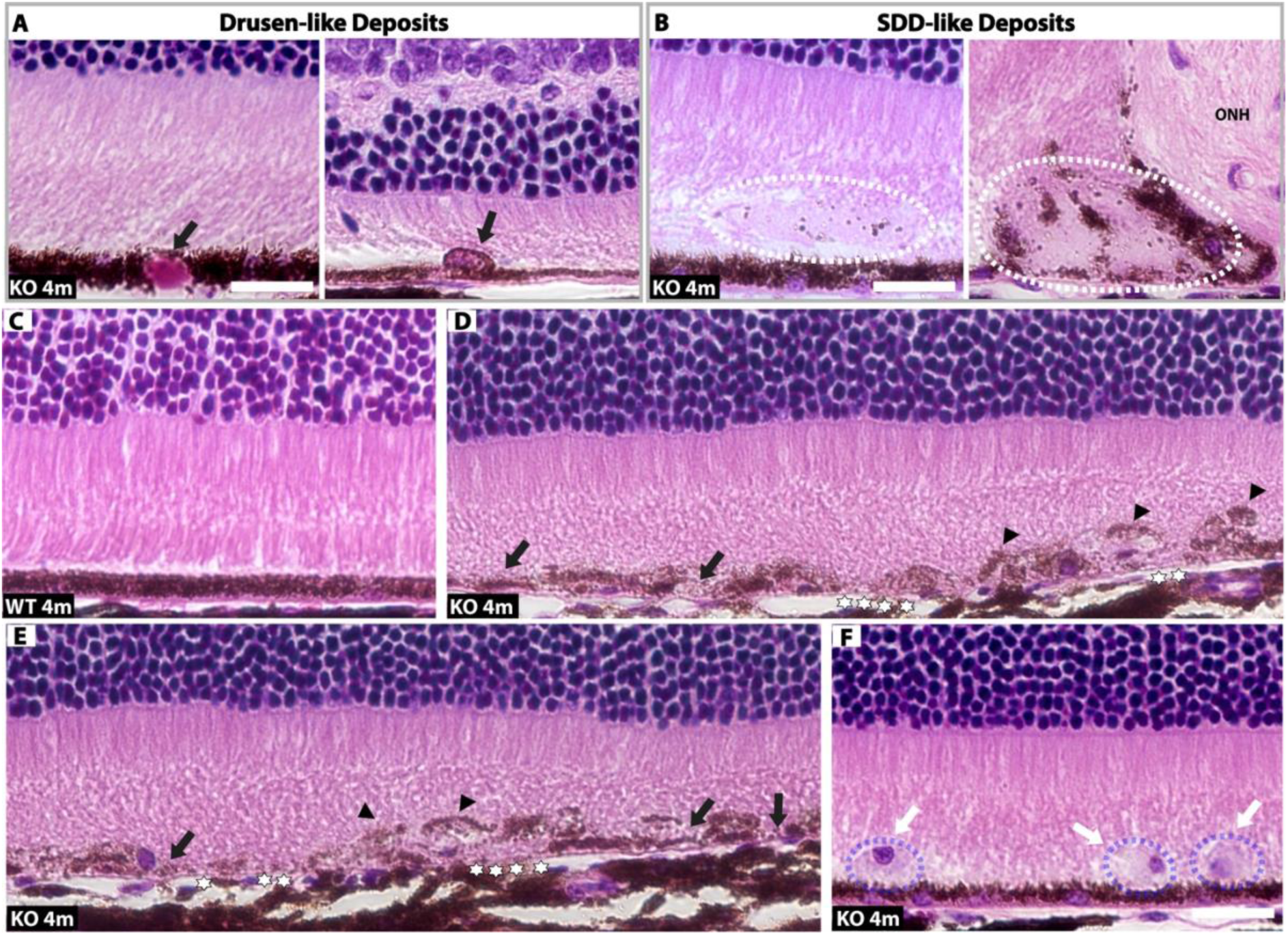
Hematoxylin- and Eosin-stained retinal sections from *Prcd*^-/-^ mice show drusenoid deposits and RPE defects. A. Representative images from 4-month-old *Prcd^-/-^*mice showing drusen-like focal deposits (*black arrows*). **B.** H&E images showing SDD-like focal deposits (*dashed ovals*) in the subretinal space. **C**. Representative image from 4-month-old WT showing normal RPE morphology. **D, E.** Serial H&E images of *Prcd^-/-^* retina showing RPE atrophy and thinning (*arrows*), RPE-free zones (*stars*) and RPE sloughing (*arrowheads*). **E.** H&E image showing infiltration of non-pigmented cells (*arrows and dashed purple ovals*) in the subretinal space of *Prcd^-/-^* retina. ONH – optic nerve head. Scale bars – 20µm.

To investigate whether the focal deposits observed in *Prcd^-/-^* mice are drusenoid in nature, we performed immunohistochemical analysis on retinal cryosections obtained from 10 to 12-month-old *Prcd^-/-^* mice along with age-matched WT controls with known drusen associated markers –ApoE and vitronectin (VTN). While control mice showed no focal deposits, apart from the expected VTN expression in the Bruch’s membrane (BrM) (Fig. 5A) (33), *Prcd^-/-^* mice exhibited focal deposits between the RPE-BrM that stained positive for ApoE and VTN (Fig. 5B, *arrows*). SDD-like focal deposits found in *Prcd^-/-^* mice also stained positive for ApoE and VTN (Fig. 5C, *dashed ovals*). In addition, transmission electron microscopy (TEM) analyses also revealed SDD-like focal deposits (Fig. 5D, *dashed box*) that presented with membrane-enclosed structures resembling lipoproteins (Fig. 5D, *magnified inset, arrows*) and aggregates of lipoproteins (Fig. 5D, *magnified inset, dashed oval*). These data collectively suggest RPE defects in the PRCD-deficient retina. More importantly, our data demonstrate AMD-like clinicopathological features in *Prcd^-/-^*mice.

**Figure 5.**
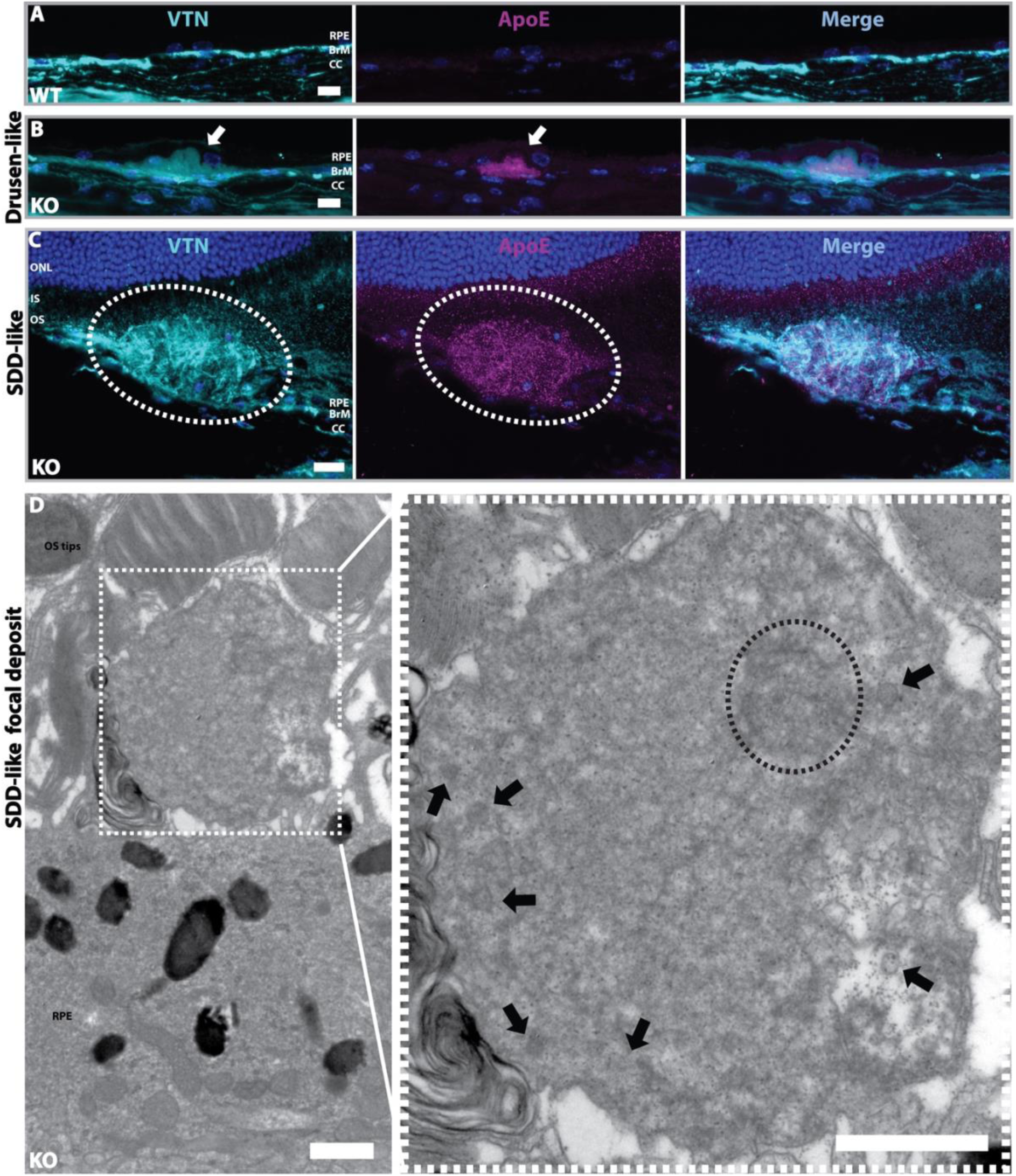
PRCD-deficient retina show drusen- and SDD-like focal deposits. A, B. Representative confocal images of RPE-BrM-Choroid from 10 to 12-month-old age-matched WT control (**A**) and *Prcd^-/-^* mice (**B**) stained for drusen-associated markers ApoE (*magenta*) and VTN (*cyan*) (n=3). White arrows in B panels point to the focal deposits resembling drusen. RPE nuclei are labeled with DAPI. **C.** Confocal images of *Prcd^-/-^* mice retinal cryosections showing SDD-like focal deposit (*delineated by the dashed-white oval*) expressing VTN and ApoE. ONL is labeled with DAPI. Scale bars - 10µm. **D.** TEM image from 12-month-old *Prcd^-/-^* mouse showing SDD-like focal deposit (*dashed square*) in the subretinal space. The magnified inset on the right panel shows membrane-enclosed structures resembling lipoproteins (*black arrows*) and aggregates of lipoproteins (*dashed oval*) within the focal deposit. Scale bars - 1µm.

### *Prcd*^-/-^ mice exhibit morphological and functional defects in the RPE cells

The presence of HRF (Fig. 3J; *arrows*), RPE-thinning, and atrophy in patches throughout the retina of *Prcd^-/-^* mice (Fig. 4D, E) prompted us to investigate the morphological and functional integrity of the RPE in the *Prcd^-/-^* mice. RPE cells assist in photoreceptor OS renewal by phagocytosing distal OS discs in a circadian manner (21,23). Previous studies in *Prcd^C2Y/C2Y^* canines and *Prcd^-/-^* mice report reduced OS renewal rates and impaired RPE phagocytosis, respectively. (5,8,25). We investigated the phagosome content of RPE cells using rhodopsin labeling on RPE flatmounts obtained from 6 to 7-month-old *Prcd^-/-^* (Fig. 6 *B)* and age-matched WT controls (Fig. 6 *A)* and observed a ∼55% decrease in the expression of rhodopsin-positive puncta in *Prcd^-/-^*mice (Fig. 6B, C). Our data further corroborates impaired phagocytosis in *Prcd^-/-^* mice, as reported earlier (25). Accumulation of lipids or lipid derivatives in the RPE were previously reported to result in RPE cell dysfunction and death in both genetic and age-related disorders like Stargardt disease and AMD, respectively (34). To investigate RPE dysfunction in *Prcd^-/-^* mice, we stained retinal cryosections from 4-month-old *Prcd^-/-^* and age-matched WT controls with Nile Red. We found accumulation of lipid deposits within the RPE of *Prcd^-/-^* mice (Fig. 6 *E*) that were absent in WT controls (Fig. 6 *D*).

**Figure 6.**
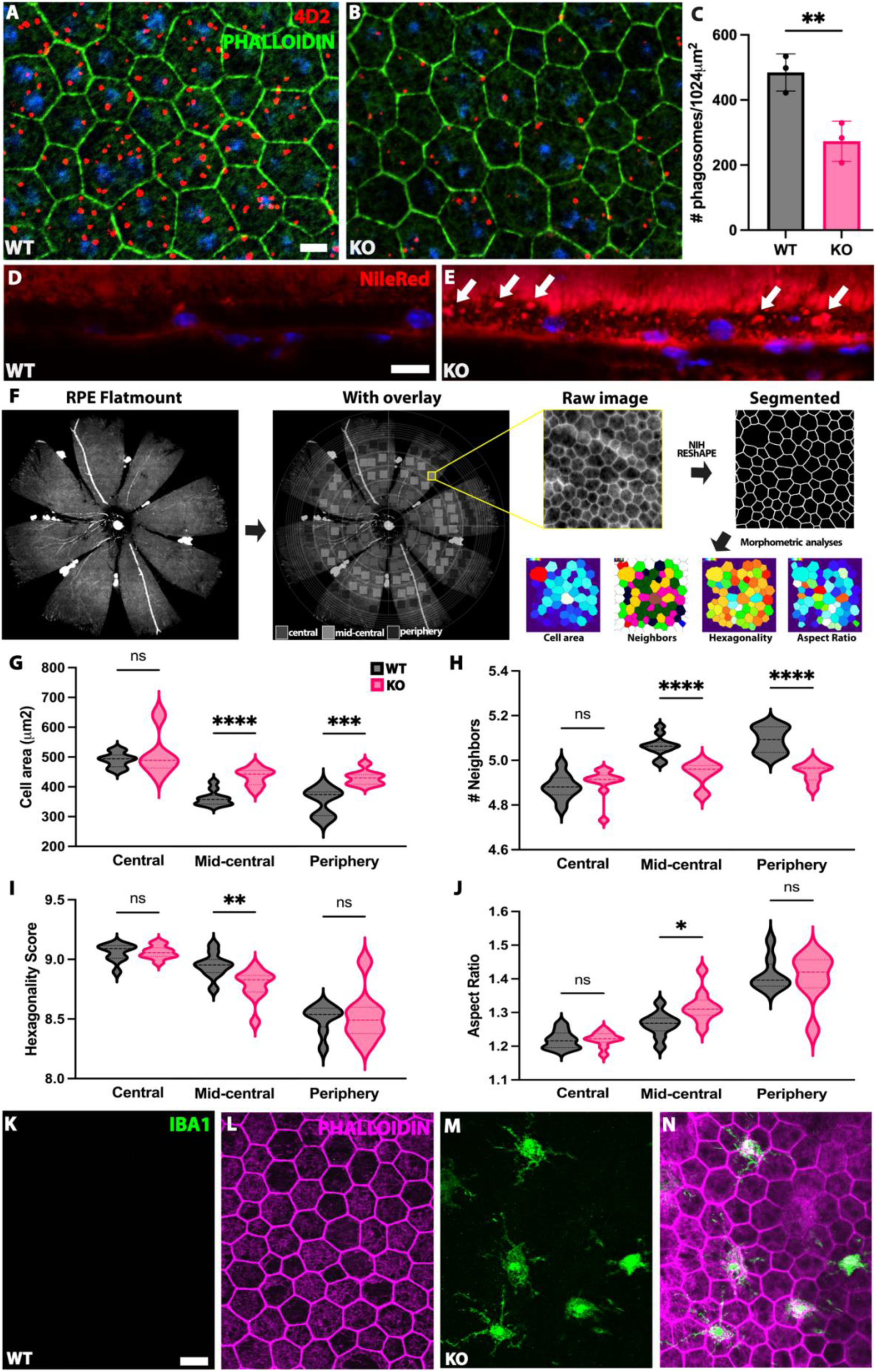
Functional and structural defects in the RPE of *Prcd*^-/-^ mice. **A, B.** Representative confocal images of RPE flatmounts from 6- to 7-month-old WT control (A) and *Prcd^-/-^*(B) mice labelled with 4D2 (*red*) and phalloidin (*green*). **C.** Quantification of the number of phagosomes per 1024µm^2^ area. Groups were compared using a two-tailed Student t-test (n=3 biological replicates, 15 technical replicates; p<0.01). Data are represented as mean ± SD. **D, E.** Nile Red (*red*) staining of Retina-RPE-choroid from 4-month-old WT controls and *Prcd^-/-^* mice showing neutral lipid accumulation (E, *arrows*) in the RPE of *Prcd*^-/-^ mice. **F.** Schematic showing RPE flatmount data sampling and morphometric analyses of RPE cells using NIH REShAPE software (35). **G-J.** Quantification of RPE cell areas (G), number of neighbors (H), hexagonality (I) and aspect ratios (J) from 12-month-old WT (*grey*) and *Prcd^-/-^* mice (*pink*). Groups were compared using unpaired two-tailed Student t-test with Welch’s correction (n=3 biological replicates, 6-10 technical replicates; over 1000 cells were examined from each region; *P<0.05; **P<0.002; ***P<0.001; ****P<0.0001). Dashed line in the center represents the median, dotted lines at the bottom and top indicate the first and third quartiles, respectively. **K-N.** RPE flatmounts from 12-month-old WT (K, L) and *Prcd^-/-^*mice (M, N) labelled with IBA1 (*green*) and Phalloidin (*magenta*) showing subretinal microglial/macrophage infiltration in *Prcd*^-/-^ mice.

To further investigate RPE health, we used phalloidin labeling on RPE flatmounts from 10 to 12-month-old *Prcd^-/-^* mice and age-matched WT controls and performed morphometric analysis using NIH REShAPE software (Fig. 6F) (35). We observed a significant increase in the RPE cell areas (Fig. 6G) and a decrease in the total number of neighbors (Fig. 6H) in the mid-central and peripheral regions of the eye in *Prcd^-/-^*mice. RPE cells in the mid-central region also displayed lower hexagonality scores (Fig. 6I) and higher aspect ratios (Fig. 6J) in *Prcd^-/-^* mice. However, no differences were observed between the groups in the above four morphometry parameters in the central region. Additionally, we observed infiltration of microglia into the subretinal space of *Prcd^-/-^*retina (Fig. 6 *M, N;* Fig. S4B) that was absent in WT controls (Fig. 6 *K, L*), which further supports RPE dysfunction and degeneration (36). These microglia are abundantly stained by ApoE (Fig. S3*D*) and exhibit an amoeboid phenotype (Fig. 6 *M,* Fig. S4B*, magnified inset*), which represents an activated state that is highly phagocytic (37). Together, these data provide evidence for both structural and functional defects in the RPE of *Prcd^-/-^* mice.

### *Prcd*^-/-^ mice exhibit lipofuscin accumulation and Bruch’s membrane deposits

To further dissect the lipid accumulation, impaired phagocytosis and various morphological defects observed in the RPE of *Prcd^-/-^* mice, we sought to evaluate the ultrastructure of the RPE and Bruch’s membrane (BrM) of 12-month-old *Prcd^-/-^* and age-matched WT controls using TEM. We found a higher accumulation of lipofuscin (Fig. 7G; *arrows*) and melanolipofuscin granules (Fig. 7G; *arrowheads*) in the RPE of *Prcd^-/-^* mice when compared to age-matched WT controls (Fig. 7F), suggesting lysosomal dysfunction. We further confirmed the defective phagocytosis in *Prcd^-/-^*mice (Fig. 6 *B,C*), as reported earlier (25). More interestingly, we observed thickening of BrM in *Prcd^-/-^* mice (Fig. 7B). These BrM deposits were sometimes found to comprise coated membrane-bound (CMB) bodies (Fig. 7Di, *arrows*) that ranged from 400-1000nm in size and contained coated vesicle-like (CVL) bodies (Fig. 7Di, *black arrowhead*) and lipoproteins (Fig. 7Di, *open arrowheads*). We also found electrolucent structures with coated membranes resembling CVL bodies in the elastic layer of BrM (Fig. 7 *Dii, arrow*). In addition, we observed accumulation of extracellular deposits, likely cellular debris generated by degenerating RPE cells, in the subretinal space of *Prcd^-/-^* mice (Fig. 7E, *dashed line*; Fig. S5, *dashed box*). These deposits sometimes contained lipofuscin (Fig. 7E *inset, arrow*), lamellar bodies (Fig. 7E, *open arrowheads)* and membranous vacuoles (Fig. S5; *arrowheads).* Thus, in addition to potential lysosomal impairment, our data provides sufficient evidence indicating that the RPE of *Prcd^-/-^* mice undergoes accelerated aging.

**Figure 7.**
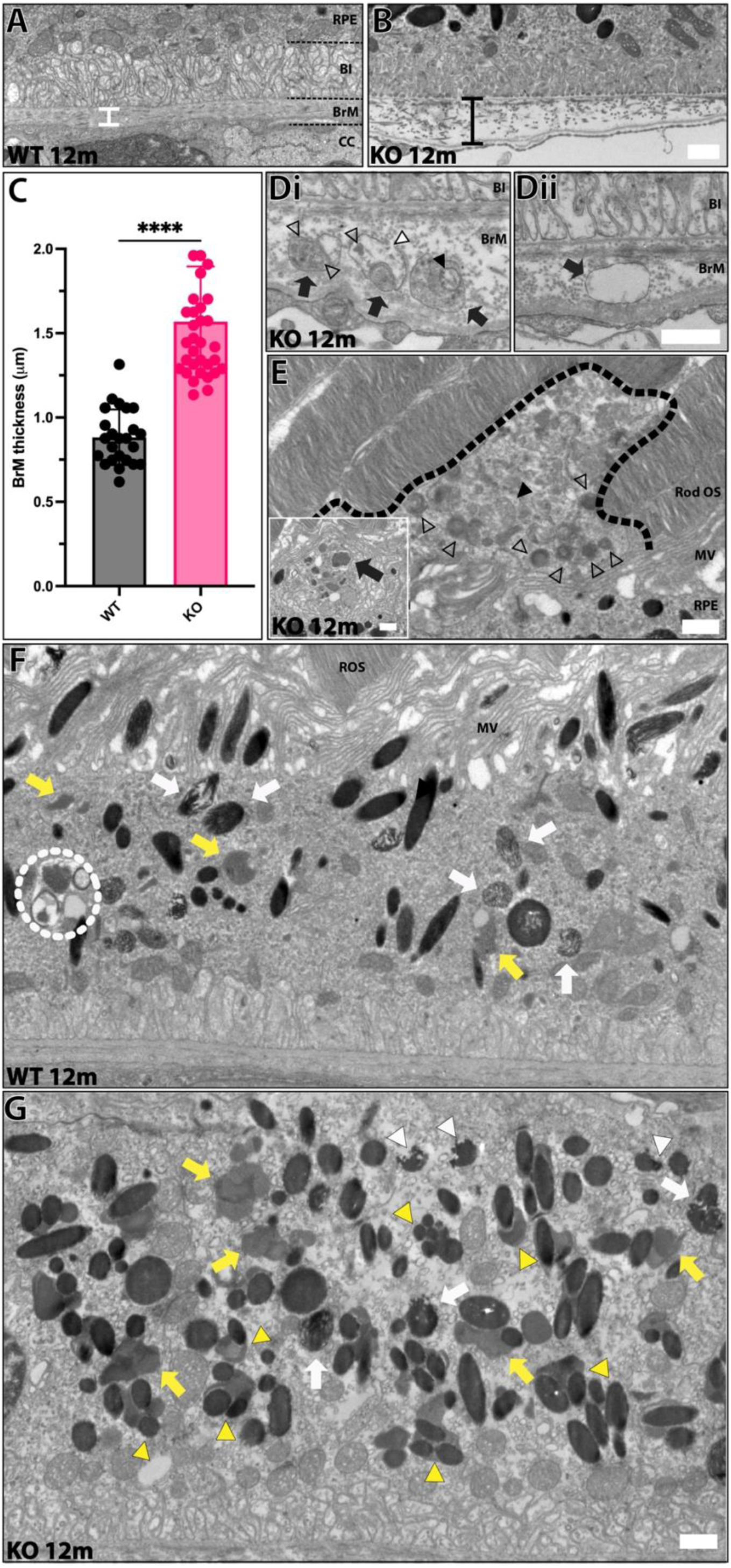
BrM thickening and lipofuscin accumulation in the RPE of *Prcd*^-/-^ mice. **A-B** Representative TEM images of BrM from 12-month-old *Prcd^-/-^* and WT mice. The black bar indicates increased BrM thickness in *Prcd*^-/-^ mice relative to the average thickness observed in WT, indicated by the white bar. **C.** Quantification of average BrM thickness in WT vs *Prcd^-/-^* mice. Mean ± SD of surveyed areas from 8-12 individual scans per genotype are shown. Groups were compared using a two-tailed Student t-test (n=3, p<0.0001). **Di.** Representative images of CMB bodies (*arrows*) containing CVL bodies (*black arrowhead*) and lipoproteins (*open arrowheads*) found within the BrM in *Prcd^-/-^* mice. White arrowhead points to a rupture in the membrane. **Dii.** Representative image of electrolucent structures found in the BrM resembling CVL bodies (*arrow*)**. E.** TEM image of 12-month-old *Prcd^-/-^*mouse showing extracellular deposits (*delineated by dashed black border*) containing lamellar bodies (*arrowheads*) in the space abutting the rod OS tips. Inset shows vesiculations within RPE encompassing structures resembling partially digested OS and lipofuscin aggregates (*arrow*). Black arrowhead points to a structure resembling autophagic cargo. **F.** RPE TEM image from 12-month-old WT mouse showing lipofuscin (*yellow arrows*), undigested OS material in phagosomes (*white arrows*) and phagolysosome (*dashed circle*) **G.** Representative TEM image from *Prcd^-/-^* mouse showing extensive accumulation of lipofuscin (*yellow arrows*), melanolipofuscin (*yellow arrowheads*) and partially digested pigment granules (*white arrowheads*). White arrows point to phagosomes. CC – choriocapillaris/choroid; BrM – Bruch’s membrane; BI – Basal infoldings; RPE – Retinal pigmental epithelium; MV – Microvilli; Rod OS – Rod outer segment. Scale bars -1µm.

## Discussion

PRCD has long been associated with RP but its precise function in maintaining photoreceptor OS structure and function remains obscure. Recent studies demonstrate the importance of PRCD in high-fidelity disc morphogenesis. However, it is unclear why nascent discs do not flatten properly in the absence of PRCD (7). Distinct differences in the total fatty acid composition, and cholesterol to phospholipid ratios between the rod OS disc and plasma membranes (9,10), suggest that a tight regulation of membrane lipid composition is imperative for proper disc structure and function. In this study, we investigated potential alterations in retinal cholesterol levels, including phospholipid composition in *Prcd^-/-^* mice. We show altered levels of total retinal cholesterol in these mice. Our findings support the accumulation of cholesteryl esters (C.Es) and lipid deposits in the photoreceptors and the RPE of *Prcd^-/-^*mice. More interestingly, we found drusen- and SDD-like focal deposits in addition to extensive lipofuscin accumulation in our *Prcd^-/-^* mice, indicating an AMD-like phenotype.

Cholesterol favors a saturated over unsaturated fatty acyl environment (16,38,39). Rod OS discs get increasingly enriched in DHA as they are apically displaced, progressively increasing disc unsaturation which promotes cholesterol depletion (40,41). Reduced rod OS DHA levels reported in canine *Prcd^C2Y/C2Y^* models (17) imply a higher-than-normal degree of disc saturation which can both retain and accumulate rod OS membrane cholesterol. It is well known that excessive retinal cholesterol in the photoreceptor OS is esterified (42–44). Our untargeted lipidomic analyses of retinal tissue show accumulation of C.Es in addition to decreased DHA levels in *Prcd^-/-^* mice. Furthermore, increased expression of ADRP/PLIN2 in the photoreceptor IS and OS in *Prcd^-/-^* mice also supports elevated C.E levels. Overexpression of ADRP (PLIN1 and PLIN2) proteins has been previously shown to promote lipid droplet (LD) accumulation in drosophila photoreceptors (45). Thus, the excess retinal cholesterol in *Prcd*^-/-^ mice may be esterified to be stored in LDs.

Membrane lipid composition can modulate bilayer properties and rhodopsin activation (46–48). Attenuated scotopic responses were previously reported in *Prcd^-/-^* mice (6,25). We propose that the high cholesterol levels in *Prcd^-/-^* mice can lead to increased attenuation of light-induced conformational changes and activation of rhodopsin, either by further reducing the partial free volume of the bilayer or through direct interaction with rhodopsin, or both (49–51). Additionally, the decreased levels of DHA could also contribute to the attenuated visual responses in *Prcd^-/-^* mice. DHA also interacts with rhodopsin, but, on the contrary, it increases the membrane fluidity and augments the kinetics of photocycle by stabilizing the active state of rhodopsin (52–55). Also, as cholesterol can increase the membrane bending rigidity of the phospholipid bilayer (56–58), it is interesting to speculate whether the high cholesterol and low DHA levels in the absence of PRCD, can lead to an increased overall membrane bending rigidity of the rod OS discs. This could perhaps, at least in part, explain the disc flattening issue observed in *Prcd^-/-^*mice (7). Moreover, high cholesterol could also alter the nanodomain organization of rhodopsin in the disc membranes, as the preferential interaction of cholesterol with saturated lipids promotes lateral separation of rhodopsin into membrane areas enriched with DHA or unsaturated fatty acids (59). This notion is supported by the altered rhodopsin distribution and packaging densities that we previously reported using our *Prcd^-/-^* mouse model (6).

RPE is a monolayer of polarized, post-mitotic cells that forms the blood-retina barrier and performs multiple functions critical for the visual cycle and retinal homeostasis (60,61). Several RP models demonstrate structural alterations and atrophy in the RPE that present as various fundus abnormalities, including bone-spicule pigmentation, the hallmark feature of RP, among others (62). Intriguingly, *Prcd^-/-^* mice developed hypo-pigmentary fundus lesions as early as 2 months of postnatal age that were virtually absent from age-matched wildtype controls. These hypo-pigmentary changes could result from focal RPE atrophy or degeneration, which is supported by – (a) RPE thinning and RPE-free zones as observed in H&E analysis, (b) RPE cellular debris in the subretinal space as observed in TEM analyses, and (c) Morphological degenerative changes in the RPE of *Prcd^-/-^* mice as demonstrated by RPE morphometric analyses. We did not observe any pigmentation resembling bone spicules in our model, perhaps because the photoreceptors remain viable despite the observed progressive structural and functional deficits. A previous study on rhodopsin knockout mice demonstrated that bone spicules only form in the areas that completely lack photoreceptors, as it allows a direct contact between the RPE and inner retinal vessels, which triggers RPE migration (63).

Various retinal dystrophies, whether inherited or age-related, present with accumulation of lipid deposits within the RPE and BrM (33,64,65). We show accumulation of lipid deposits in the RPE of *Prcd^-/-^* mice. More importantly, we report sub-RPE and sub-retinal focal deposits in *Prcd^-/-^* retina that resembled drusen and SDD in their composition, as demonstrated by the expression of drusen-associated markers - ApoE and vitronectin (66–68). It is widely known that both drusen and SDD are risk factors for AMD (69–71). Cholesterol homeostasis has long been implicated in the pathogenesis of AMD. While drusen contains both FC and C.Es; SDD mostly contains FC (68,72). We propose that increased C.E levels may lead to the accumulation of focal deposits resembling drusen and SDD in *Prcd^-/-^* mice. Moreover, cellular debris from degenerating RPE can also promote the formation of drusenoid focal deposits (73,74). Ours is the first study to report AMD-like pathological features in *Prcd*^-/-^ mice. It is worth noting that *Prcd* is located within the 17q25 genetic locus in humans (75,76), which has been identified as one of the susceptibility loci of AMD (77).

One of the key functions of RPE is the daily phagocytosis of photoreceptor distal ends and recycling of DHA which assists in the continuous renewal of the photoreceptor OS (20–23). Impaired phagocytosis has been previously reported in *Prcd^-/-^* mice (25). In this study, we provide additional evidence of defective phagocytosis by reporting prominent accumulation of lipofuscin in the RPE of *Prcd^-/-^*mice; which can lead to lysosomal dysfunction (78). Past studies have shown that lipofuscin accumulation leads to lysosomal membrane permeabilization, eventually resulting in autophagic dysregulation that severely reduces the phagocytic capacity of the RPE and leads to RPE apoptosis (79,80). Furthermore, we show BrM deposits in *Prcd^-/-^* mice resembling BrM thickening reported in AMD, which primarily results from lipid and protein accumulation and precedes basal deposit and drusen formation (33,81,82). We also found various inclusions within these BrM deposits that likely indicate transcytosis of undigested OS material or damaged organelles by the RPE due to exceeded lysosomal capacity as reported in earlier studies (83–85). Besides, BrM deposits have been previously reported to result from inhibition of lysosomal degradation (86), further suggesting lysosomal impairment in the RPE of *Prcd^-/-^* mice. Several studies report lipofuscin accumulation, BrM and drusenoid deposits as typical features of aged RPE (81,82,87–89) suggesting accelerated aging of the RPE in *Prcd^-/-^* mice.

Although PRCD is exclusively expressed in the photoreceptor OS (1,4), our study demonstrates an array of structural and functional defects in the RPE of *Prcd*^-/-^ mice. As the RPE and photoreceptors function as a metabolic ecosystem (90,91), we conjecture that the lipid alterations observed in *Prcd*^-/-^ mice can dysregulate RPE homeostasis, leading to the observed RPE defects. For instance, the low retinal DHA (17) and impaired phagocytosis (25) reported in *Prcd*^-/-^ mice suggest a reduced free DHA pool in the RPE. This could lead to increased oxidative burden and eventual RPE dysfunction, as free DHA generated from phagocytosed OS attenuates oxidative stress-induced apoptosis in the RPE (28,92). Additionally, lipofuscin accumulation due to impaired phagocytosis can prevent cholesterol efflux in the RPE, resulting in the accumulation of both FC and C.Es (93) and lead to drusen- and SDD-like deposits observed in *Prcd*^-/-^ mice. This can in-turn exacerbate photoreceptor dysfunction and degeneration. Overall, our study provides further insights into lipid abnormalities and RPE impairment accompanying PRCD deficiency and suggests that RPE dysfunction likely contributes to photoreceptor dysfunction and degeneration in *Prcd*^-/-^ mice.

## Experimental procedures

### Animals

Mice used in this study were maintained under standard 12-hr light and 12-hr dark cycle and fed ad libitum. *Prcd*^-/-^ mice were periodically backcrossed with wildtype 129/SV-E mice to maintain the genetic integrity. Generation of global *Prcd*^-/-^ mice is described in earlier study (6). Experimental animals were genotyped by direct sequencing of polymerase chain reaction product obtained from the genomic DNA. All experimental procedures conducted were approved by the Institutional Animal Care and Use Committee of West Virginia University (IACUC Protocol # 1603001820).

### Lipid Staining

Retinal cryosections were rehydrated in 1X phosphate buffered saline (1X PBS) for 20 minutes at room temperature (RT) followed by incubation in 1.75 µg/mL Nile Red (Sigma, Cat# N3013) in 100% acetone for 1-hour at RT in the dark. Sections were washed thrice with 1X PBS + DAPI for 5 minutes each and mounted in Prolong Gold antifade reagent (Life technologies, Cat# P36934) and scanned using Nikon Eclipse Ti laser scanning confocal microscope with C2 camera (Nikon Instruments, Melville, NY).

### Immunohistochemistry

Eyes were enucleated post carbon dioxide (CO2) asphyxiation, nicked at the limbus using a 22- gauge needle and post-fixed in 4% paraformaldehyde (4% PFA) at RT for 5 minutes. Corneas were removed and the resulting eyecups were further incubated in 4% PFA for an hour under gentle agitation at RT. Eyecups were washed thrice with 1X PBS for 10 minutes each and dehydrated in 20% sucrose in 1X PBS at 4 °C overnight. After dehydration, eyecups were incubated in 1:1 mixture of 20% sucrose and Tissue-Tek optical cutting temperature (OCT; Sakura Finetek, Torrance, CA) for one hour at RT before embedding in OCT. 12-16 μm thick retinal cryosections were obtained using Leica CM1850 cryostat (Leica Biosystems, Nussloch, Germany) and mounted on Superfrost Plus slides (Fisher Scientific, Cat# 1255015). Sections were blocked with block buffer (10% normal goat serum, 0.5% triton-X 100, and 0.05% sodium azide in 1X PBS) for an hour at RT followed by incubation with primary antibodies diluted in antibody dilution buffer (5% normal goat serum, 0.5% triton-X 100, and 0.05% sodium azide in 1X PBS) at 4 °C overnight. Sections were washed with 1X PBS containing 0.1% triton-X 100 (1X PBST) four times, 15 minutes each, before incubating with appropriate Alexa flour 488/546/680 conjugated secondaries and nuclear stain DAPI for 2-3 hours at RT. Sections were washed with 1X PBST as before and mounted in Prolong Gold antifade reagent (Life technologies, Cat# P36934) and coverslipped. Confocal imaging was performed on Nikon Eclipse Ti laser scanning confocal microscope with C2 camera or Nikon AX confocal system (Nikon Instruments, Melville, NY). Identical exposure and gain settings were used for both experimental and control sections for each antibody. All images represent maximum intensity z-projections generated using ImageJ (94) or FIJI (95) with the Bio-Formats plugin (https://imagej.net/Bio-Formats).

### ADRP puncta quantifications

Quantifications of ADRP/PLIN2 puncta in the photoreceptor IS and OS were performed in Image J using Phansalkar thresholding and 0-1 circularity. Separate macros were made and used to quantify the expression of ADRP puncta in the IS and the OS. (Macros only differed in the sampling box area loaded for puncta quantification). Number of technical replicates is listed in the figure legends. T-tests were performed using GraphPad Prism Version 10.2 (La Jolla, CA). P values of <0.05 were considered significant.

### Antibodies and Dyes

All antibodies and dyes used in this study are listed in SI Appendix, Table 1

### RPE Phagosome staining and quantification

RPE flatmounts were made using the protocol described previously (96), with a few modifications. Briefly, eyes were enucleated at ZT6 and rinsed with ice-cold 1X PBS followed by fixation in 2% PFA for 2 hours at RT. Post removing cornea, iris and lens, 8 radial cuts were made to flatten the eyecup. After removing the retinal tissue, the remaining RPE/choroid/sclera were washed several times with ice-cold 1XPBS and blocked with 5% normal goat serum and 0.5% triton-X 100 in 1X PBS for 1-2 hours at RT. After blocking, RPE flatmounts were incubated with 4D2 antibody and phallodin 488 for 2 hours at RT. Post washing with 1XPBST four times, 15 minutes each, flatmounts were incubated with Alexa flour anti-mouse 546 conjugated secondary and nuclear stain DAPI for 1 hour at RT. Post 1X PBST washes as before, RPE flatmounts were mounted in Prolong Gold antifade reagent and scanned using 40X objective at 2X ZOOM on Nikon AX confocal system (Nikon Instruments, Melville, NY). Phagosome puncta quantification was performed in Image J using Phansalkar thresholding and 0-1 circularity. Number of technical replicates is listed in the figure legends. T-tests were performed using GraphPad Prism Version 10.2 (La Jolla, CA). P values of <0.05 were considered significant.

### RPE morphometric analysis

RPE flatmounts from *Prcd*^-/-^ mice and age-matched WT controls were incubated with Phalloidin 680 (Cayman chemicals, Cat#20556) and nuclear stain DAPI for 2 hours at RT. Post washes with 1X PBST as before, the flatmounts were scanned using 20X objective on Slide Scanner (Olympus, VS120). RPE flatmount images were overlayed with a grid made of concentric circles placed 320 µm apart as described before (97) and were arbitrarily partitioned into central (400 to 1040 µm distance from optic nerve), mid-central (1040-1680 µm) and peripheral regions (>1680 µm). Sampling boxes measuring 200x200 µm were placed within each region, avoiding regions with dim staining issues or missing tissue. Approximately fifty 200x200 µm boxes were sampled from each RPE flatmount image. NIH REShAPE software (https://github.com/nih-nei/REShAPE) was used to perform RPE morphometric analysis as described in Ortolan et at 2022 (35). Only the output from images where all the cells (excluding the cells on the image edges) were faithfully detected was used for the downstream analysis. Over 1000 cells were examined from each region and T-tests were performed using GraphPad Prism Version 10.2 (La Jolla, CA). P values of <0.05 were considered significant.

### Fundoscopy and Fluorescein Angiography (FA)

Mouse pupils were dilated with Tropi-Phen ophthalmic solution (Pine Pharmaceuticals) containing 1% Tropicamide and 2.5% phenylephrine HCL for 5 minutes and later anesthetized by an intramuscular injection of ketamine (80 mg/kg) and Xylazine (10 mg/kg) at a dosage of 0.1 mL/20g body weight. GenTeal tears lubricant eye gel was liberally used throughout the procedure to prevent corneal desiccation. Regular fundus images were obtained from both eyes using MICRON IV ophthalmic imaging system (Phoenix-Micron Inc, OR, USA). For FA, 10% fluorescein in 1XPBS was injected intraperitoneally and the retinal vascular was imaged 5 minutes and 10 minutes post injection. Mice were administered Anti-Sedan (Orion Corporation) intraperitoneally to reverse the anesthesia immediately post procedure. Eyes were treated with Neomycin and Polymyxin B Sulfates and Bacitracin Zinc Ophthalmic ointment (Bausch & Lomb, Valeant Pharmaceuticals) to treat/prevent possible eye infections. Mice were returned to warm pad and monitored during recovery.

### Spectral Domain Optical Coherence Tomography and reflectivity quantification

Mouse pupils were dilated for a minimum of 5 minutes and later mice were anesthetized as before. *In vivo* cross-sectional images of the retina were obtained using Bioptigen R-series spectral domain ophthalmic imaging system (Bioptigen, Inc., Durham, NC). Systane gel eye drops were used to prevent corneal desiccation throughout the procedure. Longitudinal reflectance profiles (LRPs) were obtained using ImageJ as previously described with some modifications (98). Briefly, averaged images from WT and KO mice were rotated 90 degrees and LRPs were obtained halfway from the optic nerve to image edge using the line tool in Image J set to 50-pixel horizontal width. The ROIs for LRPs are positioned starting from the ONL band to RPE band. The resultant averaged LRPs were aligned and normalized as previously described (98).

### Lipid Extraction, Untargeted LC/MS and Data Analysis

Mice were euthanized by cervical dislocation to avoid any confounding effects of CO2 inhalation on the lipid composition. Immediately after enucleation, retinas were dissected and flash-frozen in liquid nitrogen until further processing. Untargeted lipidomic profiling, combined with lipid identification and quantitation was conducted by Creative Proteomics Inc. (NY, USA). Briefly, retinal samples were homogenized in 75% methanol containing 1mM Butylated Hydroxytoluene (BHT) using zirconium beads and bullet blender tissue homogenizer. Retinal homogenates were spiked with 10µL of internal standard consisting of 100µM di-myristoyl phospholipids (PG, PE, PS, PA, PC (46:0), SM (30:1) and 25 µM TG (14:1). To extract lipids, 1mL of methyl-tert-butyl ether (MTBE) and 60 µL methanol were added to samples and vortexed for 60 min at RT, followed by addition of 170 µL water and vortexing for 15 minutes. Supernatants were collected post 15-minute centrifugation and protein pellets were reextracted as above and pooled before drying in a speedvac. Dried lipids were resuspended in 0.01% BHT and were diluted in isopropanol:methanol (2:1, v:v) containing 20mM ammonium formate immediately prior to the analyses. Shimadzu Prominance HPLC was used to deliver 5µL sample directly by flow injection at a rate of 10μL/min. Thermo Scientific LTQ-Orbitrap Velos mass spectrometer was used to obtain Full scan MS spectra at 100,000 resolution (defined at m/z 400) in both positive and negative ionization modes. Offline mass recalibration was done using Thermo Xcalibur software according to the vendor’s instructions. Linear fit algorithm in Lipid Mass Spectrum Analysis (LIMSA) v.1.0 software was used to identify lipids. Peaks were found and corrected for 13°C isotope effects using in-house database of hypothetical lipid compounds and peak areas are quantified by normalization against internal standards. Xcalibur software was used to perform structural analysis on top ∼300 most abundant peaks in both positive and negative ionization modes. The relative abundance estimates of various lipid species are reported in SI Appendix, Table S1. Data analysis was performed using GraphPad prism software. Groups were compared using unpaired multiple t-tests with assumptions of equal SDs. No corrections for multiple comparisons were made.

### H&E staining

Eyes were enucleated post CO2 asphyxiation and fixed overnight in 1mL of Excalibur Alcoholic Z-fix (Excalibur pathology Inc., Norman, OK) at RT. Post embedding in paraffin, 4µm-thick serial sections through the entire eye were obtained and the standard Hematoxylin-Eosin staining was performed. Slides were scanned using Nikon inverted brightfield Eclipse Ti microscope with DS-Ri2 camera (Nikon Instruments, Melville, NY).

### Transmission Electron Microscopy

Eyes were enucleated post CO2 asphyxiation and rinsed with milliQ water to remove blood. Next, the eyes were immersed in a petri dish with fixative solution containing 2% paraformaldehyde (EMS, Cat# 15700) + 2.5% glutaraldehyde (EMS, Cat#16020) in 100mM cacodylate buffer (EMS, Cat# 11653), (pH 7.4). A small incision is made on the Ora serrata, and the eyes were fixed for 1-hour, undisturbed at RT. After removing the cornea, iris and lens, the eyecups were returned to the above fixative solution in half dram vials and incubated for 2 days at RT, under gentle rolling. Post fixation, the eyecups were cut into trapezoid/rectangular shaped pieces and rinsed with 100 mM cacodylate buffer 5 times at RT. Tissues were incubated in 2% Osmium tetroxide (EMS, Cat# 19170) in 200 mM cacodylate buffer for 1-hour on ice, and transferred to 1% Uranyl acetate (EMS, Cat# 22400) and incubated overnight at 4°C. Post 2-3 rinses in milliQ water, tissue sections were then dehydrated in an ethanol series: 30%-50%-70%-95% (diluted in water); for 15 minutes each at RT. Next, the sections were dehydrated two times in 100% ethanol, for 15 minutes each, under gentle rolling. Post washing once in propylene oxide (EMS, Cat#20401) for 10 minutes under gentle rolling at RT, the sections were then transferred to a 3:1 mixture of propylene oxide: Araldite resin (EMS, Cat#13940) for overnight incubation under gentle rolling at RT. Lastly, the sections were incubated twice in 100% Araldite resin for 2 hours each at RT before embedding in fresh resin and cured at 60°C for 24-48 hours. Post trimming the extra resin, 70 nm sections were cut using Diatome Ultra 45° diamond knife and transferred onto square 100 mesh copper grids (EMS, Cat# G100-Cu). Grids were post-stained in Reynold’s lead citrate stain (EMS, Cat# 22410-01) for 2 minutes and rinsed with milliQ water 6 times and dried overnight. Post-stained sections were scanned using JEOL JEM-1010 transmission electron microscope (JEOL, Peabody, MA) equipped with Hamamatsu Orcs-HR digital camera. BrM thickness quantifications were performed using the line tool in Image J. Number of technical replicates is listed in the figure legends. T-tests were performed using GraphPad Prism Version 10.2 (La Jolla, CA). P values of <0.05 were considered significant.

## Data availability

Lipidomics data are available from the corresponding author upon request. All other data are contained within the manuscript.

## Supporting information

This article contains supporting information.

## Acknowledgments

The authors are very grateful to Dr. Christine Curcio for critical feedback and guidance on TEM experiments. We thank Drs. Visvanathan Ramamurthy, Wen Tao, Maxim Sokolov, Peter Mathers, Peter Stoilov, and Roberta Leonardi for helpful discussions and feedback. We thank Dr. Michael Robichaux and Dr. Neil Billington for technical support and guidance on image acquisition and analysis. We also thank Thamaraiselvi Saravanan and Siyan Zhu for excellent technical assistance.

## Ethics

Animal Experiments: This study was performed in strict accordance with the recommendations in the guide for the Care and Use of Laboratory Animals of the National Institute of Health. Animal studies were carried out under approved protocol from animal studies guidelines at West Virginia University protocols (IACUC #1603001820)

## Abbreviations

PRCD: Progressive rod-cone degeneration
RP: Retinitis pigmentosa
OS: Outer segment
AMD: Age-related macular degeneration
BrM: Bruch’s membrane
C.Es: Cholesteryl esters
PC: Phosphatidyl choline
PE: Phosphatidyl ethanolamine
SM: Sphingomyelin
DHA: Docosahexaenoic acid
ADRP/PLIN2: Adipocyte differentiation-related protein/perilipin 2
LD: Lipid droplet
SD-OCT: Spectral-domain optical coherence tomography
HRF: Hyperreflective foci
ApoE: Apolipoprotein E
VTN: Vitronectin
SDD: Subretinal drusenoid deposits
CVL: Coated vesicle-like
CMB: Coated membrane-bound
TEM: Transmission electron microscopy

## References

1. Skiba, N. P., Spencer, W. J., Salinas, R. Y., Lieu, E. C., Thompson, J. W., and Arshavsky, V. Y. (2013) Proteomic identification of unique photoreceptor disc components reveals the presence of PRCD, a protein linked to retinal degeneration. Journal of Proteome Research 12, 3010–3018

2. Hamel, C. (2006) Retinitis pigmentosa. Orphanet Journal of Rare Diseases 1, 1–12

3. Zangerl, B., Goldstein, O., Philp, A. R., Lindauer, S. J. P., Pearce-Kelling, S. E., Mullins, R. F., Graphodatsky, A. S., Ripoll, D., Felix, J. S., Stone, E. M., Acland, G. M., and Aguirre, G. D. (2006) Identical Mutation in a Novel Retinal Gene Causes Progressive Rod-Cone Degeneration (prcd) in Dogs and Retinitis Pigmentosa in Man. Genomics 88, 551–551

4. Murphy, J., and Kolandaivelu, S. (2016) Palmitoylation of Progressive Rod-Cone Degeneration (PRCD) Regulates Protein Stability and Localization. The Journal of Biological Chemistry 291, 23036–23036

5. Aguirre, G., Alligood, J., O’Brien, P., and Buyukmihci, N. (1982) Pathogenesis of progressive rod-cone degeneration in miniature poodles. Investigative Ophthalmology and Visual Science 23, 610–630

6. Sechrest, E. R., Murphy, J., Senapati, S., Goldberg, A. F. X., Park, P. S. H., and Kolandaivelu, S. (2020) Loss of PRCD alters number and packaging density of rhodopsin in rod photoreceptor disc membranes. Scientific Reports 2020 10:1 10, 1–12

7. Spencer, W. J., Ding, J. D., Lewis, T. R., Yu, C., Phan, S., Pearring, J. N., Kim, K. Y., Thor, A., Mathew, R., Kalnitsky, J., Hao, Y., Travis, A. M., Biswas, S. K., Lo, W. K., Besharse, J. C., Ellisman, M. H., Saban, D. R., Burns, M. E., and Arshavsky, V. Y. (2019) PRCD is essential for high-fidelity photoreceptor disc formation. Proceedings of the National Academy of Sciences of the United States of America 116, 13087–13096

8. Aguirre, G., and O’Brien, P. (1986) Morphological and biochemical studies of canine progressive rod-cone degeneration. 3H-fucose autoradiography. Investigative Ophthalmology & Visual Science 27, 635–655

9. Boesze-Battaglia, K., and Albert, A. D. (1992) Phospholipid distribution among bovine rod outer segment plasma membrane and disk membranes. Experimental eye research 54, 821–823

10. Boesze-Battaglia, K., Hennessey, T., and Albert, A. D. (1989) Cholesterol heterogeneity in bovine rod outer segment disk membranes. Journal of Biological Chemistry 264, 8151–8155

11. Fliesler, A. J., and Anderson, R. E. (1983) Chemistry and metabolism of lipids in the vertebrate retina. Progress in Lipid Research 22, 79–131

12. Daemen, F. J. M. (1973) Vertebrate rod outer segment membranes. Biochimica et Biophysica Acta (BBA) - Reviews on Biomembranes 300, 255–288

13. Huber, T., Rajamoorthi, K., Kurze, V. F., Beyer, K., and Brown, M. F. (2002) Structure of docosahexaenoic acid-containing phospholipid bilayers as studied by 2H NMR and molecular dynamics simulations. Journal of the American Chemical Society 124, 298–309

14. 14. Rodriguez De Turco, E. B., Jackson, F. R., Parkins, N., and Gordon, W. C. (2000) Strong association of unesterified [3H]docosahexaenoic acid and [3H-docosahexaenoyl]phosphatidate to rhodopsin during in vivo labeling of frog retinal rod outer segments. Neurochemical Research 25, 695–703

15. Stinson, A. M., Wiegand, R. D., and Anderson, R. E. (1991) Fatty acid and molecular species compositions of phospholipids and diacylglycerols from rat retinal membranes. Experimental eye research 52, 213–218

16. House, K., Badgett, D., and Albert, A. D. (1989) Cholesterol movement between bovine rod outer segment disk membranes and phospholipid vesicles. Experimental eye research 49, 561–572

17. Aguirre, G. D., Acland, G. M., Maude, M. B., and Anderson, R. E. (1997) Diets enriched in docosahexaenoic acid fail to correct progressive rod-cone degeneration (prcd) phenotype. Investigative Ophthalmology and Visual Science 38, 2387–2407

18. 18. Anderson, R. E., Maude, M. B., and Bok, D. (2001) Low docosahexaenoic acid levels in rod outer segment membranes of mice with rds/peripherin and P216L peripherin mutations. in Investigative Ophthalmology & Visual Science

19. Anderson, E., and Aude, M. (1991) Plasma Lipid Abnormalities in the Miniature Poodle with Progressive Rod-Cone Degeneration. Experimental Eye Research 52, 349–355

20. Chen, H., and Anderson, R. E. (1993) Metabolism in frog retinal pigment epithelium of docosahexaenoic and arachidonic acids derived from rod outer segment membranes. Experimental eye research 57, 369–377

21. LaVail, M. M. (1976) Rod outer segment disk shedding in rat retina: relationship to cyclic lighting. Science (New York, N.Y.) 194, 1071–1074

22. Young, R. W. (1967) THE RENEWAL OF PHOTORECEPTOR CELL OUTER SEGMENTS. The Journal of Cell Biology 33, 61–61

23. Young, R. W., and Bok, D. (1969) PARTICIPATION OF THE RETINAL PIGMENT EPITHELIUM IN THE ROD OUTER SEGMENT RENEWAL PROCESS. The Journal of Cell Biology 42, 392–392

24. 24. Chen, H., Ray, J., Scarpino, V., Acland, G. M., Aguirre, G. D., and Anderson, R. E. (1999) Synthesis and release of docosahexaenoic acid by the RPE cells of prcd-affected dogs. in Investigative Ophthalmology & Visual Science

25. Allon, G., Mann, I., Remez, L., Sehn, E., Rizel, L., Nevet, M. J., Perlman, I., Wolfrum, U., and Ben-Yosef, T. (2019) PRCD is concentrated at the base of photoreceptor outer segments and is involved in outer segment disc formation. Human Molecular Genetics 28, 4078–4088

26. D’Cruz, P. M., Yasumura, D., Weir, J., Matthes, M. T., Abderrahim, H., LaVail, M. M., and Vollrath, D. (2000) Mutation of the receptor tyrosine kinase gene Mertk in the retinal dystrophic RCS rat. Human molecular genetics 9, 645–651

27. Marmorstein, A. D., Johnson, A. A., Bachman, L. A., Andrews-Pfannkoch, C., Knudsen, T., Gilles, B. J., Hill, M., Gandhi, J. K., Marmorstein, L. Y., and Pulido, J. S. (2018) Mutant Best1 Expression and Impaired Phagocytosis in an iPSC Model of Autosomal Recessive Bestrophinopathy. Scientific Reports 2018 8:1 8, 1-14

28. Mukherjee, P. K., Marcheselli, V. L., Vaccari, J. C. D. R., Gordon, W. C., Jackson, F. E., and Bazan, N. G. (2007) Photoreceptor outer segment phagocytosis attenuates oxidative stress-induced apoptosis with concomitant neuroprotectin D1 synthesis. Proceedings of the National Academy of Sciences of the United States of America 104, 13158–13158

29. Elsey, D., Jameson, D., Raleigh, B., and Cooney, M. J. (2007) Fluorescent measurement of microalgal neutral lipids. Journal of Microbiological Methods 68, 639–642

30. Brasaemle, D. L., Barber, T., Wolins, N. E., Serrero, G., Blanchette-Mackie, E. J., and Londos, C. (1997) Adipose differentiation-related protein is an ubiquitously expressed lipid storage droplet-associated protein. Journal of lipid research 38, 2249–2263

31. Bartz, R., Li, W. H., Venables, B., Zehmer, J. K., Roth, M. R., Welti, R., Anderson, R. G. W., Liu, P., and Chapman, K. D. (2007) Lipidomics reveals that adiposomes store ether lipids and mediate phospholipid traffic. Journal of lipid research 48, 837–847

32. Olzmann, J. A., and Carvalho, P. (2019) Dynamics and functions of lipid droplets. Nature reviews. Molecular cell biology 20, 137–137

33. Curcio, C. A., and Johnson, M. (2012) Structure, Function, and Pathology of Bruch’s Membrane. Retina Fifth Edition 1, 465–481

34. Pan, C., Banerjee, K., Lehmann, G. L., Almeida, D., Hajjar, K. A., Benedicto, I., Jiang, Z., Radu, R. A., Thompson, D. H., Rodriguez-Boulan, E., and Nociari, M. M. (2021) Lipofuscin causes atypical necroptosis through lysosomal membrane permeabilization. Proceedings of the National Academy of Sciences of the United States of America 118

35. Ortolan, D., Sharma, R., Volkov, A., Maminishkis, A., Hotaling, N. A., Huryn, L. A., Cukras, C., Marco, S. D., Bisti, S., and Bharti, K. (2022) Single-cell–resolution map of human retinal pigment epithelium helps discover subpopulations with differential disease sensitivity. Proceedings of the National Academy of Sciences of the United States of America 119, e2117553119–e2117553119

36. Combadière, C., Feumi, C., Raoul, W., Keller, N., Rodéro, M., Pézard, A., Lavalette, S., Houssier, M., Jonet, L., Picard, E., Debré, P., Sirinyan, M., Deterre, P., Ferroukhi, T., Cohen, S. Y., Chauvaud, D., Jeanny, J. C., Chemtob, S., Behar-Cohen, F., and Sennlaub, F. (2007) CX3CR1-dependent subretinal microglia cell accumulation is associated with cardinal features of age-related macular degeneration. The Journal of Clinical Investigation 117, 2920–2920

37. Langmann, T. (2007) Microglia activation in retinal degeneration. Journal of Leukocyte Biology 81, 1345–1351

38. Van Dijck, P. W. M., De Kruijff, B., Van Deenen, L. L. M., De Gier, J., and Demel, R. A. (1976) The preference of cholesterol for phosphatidylcholine in mixed phosphatidylcholine-phosphatidylethanolamine bilayers. Biochimica et Biophysica Acta (BBA) - Biomembranes 455, 576–587

39. Yeagle, P. L., and Young, J. E. (1986) Factors contributing to the distribution of cholesterol among phospholipid vesicles. Journal of Biological Chemistry 261, 8175–8181

40. Albert, A. D., Young, J. E., and Paw, Z. (1998) Phospholipid fatty acyl spatial distribution in bovine rod outer segment disk membranes. Biochimica et Biophysica Acta (BBA) - Biomembranes 1368, 52–60

41. Schultz, Z. D. (2011) Raman Spectroscopic Imaging of Cholesterol and Docosahexaenoic Acid Distribution in the Retinal Rod Outer Segment. Australian Journal of Chemistry 64, 611–616

42. Chang, T. Y., Li, B. L., Chang, C. C. Y., and Urano, Y. (2009) Acyl-coenzyme A:cholesterol acyltransferases. American Journal of Physiology - Endocrinology and Metabolism 297, E1–E1

43. Saadane, A., Mast, N., Dao, T., Ahmad, B., and Pikuleva, I. A. (2016) Retinal Hypercholesterolemia Triggers Cholesterol Accumulation and Esterification in Photoreceptor Cells. The Journal of Biological Chemistry 291, 20427–20427

44. Zaidi, S. A. H., Lemtalsi, T., Xu, Z., Santana, I., Sandow, P., Labazi, L., Caldwell, R. W., Caldwell, R. B., and Rojas, M. A. (2023) Role of acyl-coenzyme A: cholesterol transferase 1 (ACAT1) in retinal neovascularization. Journal of Neuroinflammation 20, 1–18

45. 45. Girard, V., Jollivet, F., Knittelfelder, O., Celle, M., Arsac, J. N., Chatelain, G., Van Den-Brink, D. M., Baron, T., Shevchenko, A., Kühnlein, R. P., Davoust, N., and Mollereau, B. (2021) Abnormal accumulation of lipid droplets in neurons induces the conversion of alpha-Synuclein to proteolytic resistant forms in a Drosophila model of Parkinson’s disease. PLOS Genetics 17, e1009921–e1009921

46. Gibson, N. J., and Brown, M. F. (1993) Lipid headgroup and acyl chain composition modulate the MI-MII equilibrium of rhodopsin in recombinant membranes. Biochemistry 32, 2438–2454

47. Mitchell, D. C., Niu, S. L., and Litman, B. J. (2001) Optimization of receptor-G protein coupling by bilayer lipid composition I: kinetics of rhodopsin-transducin binding. The Journal of biological chemistry 276, 42801–42806

48. Niu, S. L., Mitchell, D. C., and Litman, B. J. (2001) Optimization of receptor-G protein coupling by bilayer lipid composition II: formation of metarhodopsin II-transducin complex. The Journal of biological chemistry 276, 42807–42811

49. Albert, A. D., Young, J. E., and Yeagle, P. L. (1996) Rhodopsin-cholesterol interactions in bovine rod outer segment disk membranes. Biochimica et Biophysica Acta (BBA) - Biomembranes 1285, 47–55

50. Khelashvili, G., Grossfield, A., Feller, S. E., Pitman, M. C., and Weinstein, H. (2009) Structural and dynamic effects of cholesterol at preferred sites of interaction with rhodopsin identified from microsecond length molecular dynamics simulations. *Proteins: Structure*, Function, and Bioinformatics 76, 403–417

51. Mitchell, D. C., Litman, B. J., Straume, M., and Miller, J. L. (1990) Modulation of Metarhodopsin Formation by Cholesterol-Induced Ordering of Bilayer Lipids. Biochemistry 29, 9143–9149

52. Feller, S. E., and Gawrisch, K. (2005) Properties of docosahexaenoic-acid-containing lipids and their influence on the function of rhodopsin. Current Opinion in Structural Biology 15, 416–422

53. Grassfield, A., Feller, S. E., and Pitman, M. C. (2006) A role for direct interactions in the modulation of rhodopsin by ω-3 polyunsaturated lipids. Proceedings of the National Academy of Sciences of the United States of America 103, 4888–4888

54. Niu, S. L., Mitchell, D. C., Lim, S. Y., Wen, Z. M., Kim, H. Y., Salem, N., and Litman, B. J. (2004) Reduced G Protein-coupled Signaling Efficiency in Retinal Rod Outer Segments in Response to n-3 Fatty Acid Deficiency. Journal of Biological Chemistry 279, 31098–31104

55. Soubias, O., Teague, W. E., and Gawrisch, K. (2006) Evidence for Specificity in Lipid-Rhodopsin Interactions. Journal of Biological Chemistry 281, 33233–33241

56. Chakraborty, S., Doktorova, M., Molugu, T. R., Heberle, F. A., Scott, H. L., Dzikovski, B., Nagao, M., Stingaciu, L. R., Standaert, R. F., Barrera, F. N., Katsaras, J., Khelashvili, G., Brown, M. F., and Ashkar, R. (2020) How cholesterol stiffens unsaturated lipid membranes. Proceedings of the National Academy of Sciences of the United States of America 117, 21896–21905

57. Chen, Z., and Rand, R. P. (1997) The influence of cholesterol on phospholipid membrane curvature and bending elasticity. Biophysical Journal 73, 267–267

58. Gracià, R. S., Bezlyepkina, N., Knorr, R. L., Lipowsky, R., and Dimova, R. (2010) Effect of cholesterol on the rigidity of saturated and unsaturated membranes: fluctuation and electrodeformation analysis of giant vesicles. Soft Matter 6, 1472–1482

59. Polozova, A., and Litman, B. J. (2000) Cholesterol dependent recruitment of di22:6-PC by a G protein-coupled receptor into lateral domains. Biophysical journal 79, 2632–2643

60. R. Sparrrow, J., Hicks, D., and P. Hamel, C. (2010) The Retinal Pigment Epithelium in Health and Disease. Current molecular medicine 10, 802–802

61. Strauss, O. (2005) The retinal pigment epithelium in visual function. Physiological reviews 85, 845–881

62. Li, Z. Y., Possin, D. E., and Milam, A. H. (1995) Histopathology of Bone Spicule Pigmentation in Retinitis Pigmentosa. Ophthalmology 102, 805–816

63. 63. Jaissle, G. B., May, C. A., Van De Pavert, S. A., Wenzel, A., Claes-May, E., Giel, A., Szurman, P., Wolfrum, U., Wijnholds, J., Fisher, M. D., Humphries, P., and Seeliger, M. W. (2010) Bone spicule pigment formation in retinitis pigmentosa: insights from a mouse model. Graefe’s archive for clinical and experimental ophthalmology = Albrecht von Graefes Archiv fur klinische und experimentelle Ophthalmologie 248, 1063–1070

64. Fu, L., Garland, D., Yang, Z., Shukla, D., Rajendran, A., Pearson, E., Stone, E. M., Zhang, K., and Pierce, E. A. (2007) The R345W mutation in EFEMP1 is pathogenic and causes AMD-like deposits in mice. Human molecular genetics 16, 2411–2422

65. Tsai, Y. T., Li, Y., Ryu, J., Su, P. Y., Cheng, C. H., Wu, W. H., Li, Y. S., Quinn, P. M. J., Leong, K. W., and Tsang, S. H. (2021) Impaired cholesterol efflux in retinal pigment epithelium of individuals with juvenile macular degeneration. American journal of human genetics 108, 903–918

66. Anderson, D. H., Ozaki, S., Nealon, M., Neitz, J., Mullins, R. F., Hageman, G. S., and Johnson, L. V. (2001) Local Cellular Sources of Apolipoprotein E in the Human Retina and Retinal Pigmented Epithelium: Implications for the Process of Drusen Formation. American Journal of Ophthalmology 131, 767–781

67. Crabb, J. W., Miyagi, M., Gu, X., Shadrach, K., West, K. A., Sakaguchi, H., Kamei, M., Hasan, A., Yan, L., Rayborn, M. E., Salomon, R. G., and Hollyfield, J. G. (2002) From the Cover: Drusen proteome analysis: An approach to the etiology of age-related macular degeneration. Proceedings of the National Academy of Sciences of the United States of America 99, 14682–14682

68. Rudolf, M., Malek, G., Messinger, J. D., Clark, M. E., Wang, L., and Curcio, C. A. (2008) Sub-retinal drusenoid deposits in human retina: Organization and composition. Experimental eye research 87, 402–402

69. Pauleikhoff, D., Barondes, M. J., Minassian, D., Chisholm, I. H., and Bird, A. C. (1990) Drusen as risk factors in age-related macular disease. American journal of ophthalmology 109, 38–43

70. Sarks, J. P., Sarks, S. H., and Killingsworth, M. C. (1994) Evolution of soft drusen in age-related macular degeneration. Eye 1994 8:3 8, 269–283

71. Wang, J. J., Foran, S., Smith, W., and Mitchell, P. (2003) Risk of Age-Related Macular Degeneration in Eyes With Macular Drusen or Hyperpigmentation: The Blue Mountains Eye Study Cohort. Archives of Ophthalmology 121, 658–663

72. Curcio, C. A., Presley, J. B., Malek, G., Medeiros, N. E., Avery, D. V., and Kruth, H. S. (2005) Esterified and unesterified cholesterol in drusen and basal deposits of eyes with age-related maculopathy. Experimental Eye Research 81, 731–741

73. Anderson, D. H., Mullins, R. F., Hageman, G. S., and Johnson, L. V. (2002) A role for local inflammation in the formation of drusen in the aging eye. American Journal of Ophthalmology 134, 411–431

74. Johnson, L. V., Leitner, W. P., Staples, M. K., and Anderson, D. H. (2001) Complement activation and inflammatory processes in Drusen formation and age related macular degeneration. Experimental eye research 73, 887–896

75. Daiger, S. P., Sullivan, L. S., and Bowne, S. J. (2013) Genes and mutations causing retinitis pigmentosa. Clinical genetics 84, 132–141

76. Pach, J., Kohl, S., Gekeler, F., and Zobor, D. (2013) Identification of a novel mutation in the PRCD gene causing autosomal recessive retinitis pigmentosa in a Turkish family. Molecular Vision 19, 1350–1350

77. Weeks, D. E., Conley, Y. P., Tsai, H. J., Mah, T. S., Rosenfeld, P. J., Paul, T. O., Eller, A. W., Morse, L. S., Dailey, J. P., Ferrell, R. E., and Gorin, M. B. (2001) Age-related maculopathy: an expanded genome-wide scan with evidence of susceptibility loci within the 1q31 and 17q25 regions. American Journal of Ophthalmology 132, 682–692

78. Gómez-Sintes, R., Ledesma, M. D., and Boya, P. (2016) Lysosomal cell death mechanisms in aging. Ageing Research Reviews 32, 150–168

79. Sundelin, S., Wihlmark, D. U., Nilsson, S. E. G., and Brunk, U. T. (1998) Lipofuscin accumulation in cultured retinal pigment epithelial cells reduces their phagocytic capacity. Current Eye Research 17, 851–857

80. Xu, Y., Li, D., Su, G., and Cai, S. (2022) The effect of A2E on lysosome membrane permeability during blue light-induced human RPEs apoptosis. BMC Ophthalmology 22, 1–12

81. Curcio, C. A., Johnson, M., Huang, J. D., and Rudolf, M. (2010) Apolipoprotein B-containing lipoproteins in retinal aging and age-related macular degeneration. Journal of Lipid Research 51, 451–467

82. Huang, J. D., Presley, J. B., Chimento, M. F., Curcio, C. A., and Johnson, M. (2007) Age-related Changes in Human Macular Bruch’s Membrane as Seen by Quick-Freeze/Deep-Etch. Experimental eye research 85, 202–202

83. Hogan, M. J., and Alvarado, J. (1967) Studies on the human macula. IV. Aging changes in Bruch’s membrane. *Archives of ophthalmology (Chicago*, Ill*. :* 1960) **77**, 410-420

84. Killingsworth, M. C. (1987) Age-related components of Bruch’s membrane in the human eye. Graefe’s archive for clinical and experimental ophthalmology = Albrecht von Graefes Archiv fur klinische und experimentelle Ophthalmologie 225, 406–412

85. Krohne, T. U., Holz, F. G., and Kopitz, J. (2010) Apical-to-Basolateral Transcytosis of Photoreceptor Outer Segments Induced by Lipid Peroxidation Products in Human Retinal Pigment Epithelial Cells. Investigative Ophthalmology & Visual Science 51, 553–560

86. Peters, S., Reinthal, E., Blitgen-Heinecke, P., Bartz-Schmidt, K. U., and Schraermeyer, U. (2006) Inhibition of Lysosomal Degradation in Retinal Pigment Epithelium Cells Induces Exocytosis of Phagocytic Residual Material at the Basolateral Plasma Membrane. Ophthalmic Research 38, 83–88

87. Delori, F. C., Goger, D. G., and Dorey, C. K. (2001) Age-related accumulation and spatial distribution of lipofuscin in RPE of normal subjects. Investigative Ophthalmology & Visual Science 42, 1855–1866

88. Kennedy, C. J., Rakoczy, P. E., and Constable, I. J. (1995) Lipofuscin of the retinal pigment epithelium: A review. Eye 1995 9:6 9, 763–771

89. Wavre-Shapton, S. T., Meschede, I. P., Seabra, M. C., and Futter, C. E. (2014) Phagosome maturation during endosome interaction revealed by partial rhodopsin processing in retinal pigment epithelium. Journal of Cell Science 127, 3852–3861

90. Kanow, M. A., Giarmarco, M. M., Jankowski, C. S. R., Tsantilas, K., Engel, A. L., Du, J., Linton, J. D., Farnsworth, C. C., Sloat, S. R., Rountree, A., Sweet, I. R., Lindsay, K. J., Parker, E. D., Brockerhoff, S. E., Sadilek, M., Chao, J. R., and Hurley, J. B. (2017) Biochemical adaptations of the retina and retinal pigment epithelium support a metabolic ecosystem in the vertebrate eye. eLife 6

91. 91. Sinha, T., Naash, M. I., and Al-Ubaidi, M. R. (2020) The Symbiotic Relationship between the Neural Retina and Retinal Pigment Epithelium Is Supported by Utilizing Differential Metabolic Pathways. iScience 23

92. Mukherjee, P. K., Marcheselli, V. L., Serhan, C. N., and Bazan, N. G. (2004) Neuroprotectin D1: A docosahexaenoic acid-derived docosatriene protects human retinal pigment epithelial cells from oxidative stress. Proceedings of the National Academy of Sciences of the United States of America 101, 8491–8496

93. Lakkaraju, A., Finnemann, S. C., and Rodriguez-Boulan, E. (2007) The lipofuscin fluorophore A2E perturbs cholesterol metabolism in retinal pigment epithelial cells. Proc Natl Acad Sci U S A 104, 11026–11031

94. Schneider, C. A., Rasband, W. S., and Eliceiri, K. W. (2012) NIH Image to ImageJ: 25 years of image analysis. Nature Methods 2012 9:7 9, 671–675

95. Schindelin, J., Arganda-Carreras, I., Frise, E., Kaynig, V., Longair, M., Pietzsch, T., Preibisch, S., Rueden, C., Saalfeld, S., Schmid, B., Tinevez, J. Y., White, D. J., Hartenstein, V., Eliceiri, K., Tomancak, P., and Cardona, A. (2012) Fiji: an open-source platform for biological-image analysis. Nature Methods 2012 9:7 9, 676–682

96. Lew, D. S., Mazzoni, F., and Finnemann, S. C. (2020) Microglia Inhibition Delays Retinal Degeneration Due to MerTK Phagocytosis Receptor Deficiency. Frontiers in Immunology 11, 1463–1463

97. Boatright, J. H., Dalal, N., Chrenek, M. A., Gardner, C., Ziesel, A., Jiang, Y., Grossniklaus, H. E., and Nickerson, J. M. (2015) Methodologies for analysis of patterning in the mouse RPE sheet. Molecular Vision 21, 40–40

98. Bonilha, V. L., Bell, B. A., Rayborn, M. E., Yang, X., Kaul, C., Grossman, G. H., Samuels, I. S., Hollyfield, J. G., Xie, C., Cai, H., and Shadrach, K. G. (2015) Loss of DJ-1 elicits retinal abnormalities, visual dysfunction, and increased oxidative stress in mice. Experimental eye research 139, 22–22

