## Supplemental Figures 1-5 for "Aberrant lipid accumulation and retinal pigmental epithelium dysfunction in PRCD-deficient mice"

**List of materials included.**

I. **Figure S1:** *Prcd*$^{-/-}$ mice accumulate EVs.

II. **Figure S2:** FC and TAG levels are unchanged in *Prcd*$^{-/-}$ mice.

III. **Figure S3:** *Prcd*$^{-/-}$ mice exhibit extracellular focal deposits and non-pigmented cells in the subretinal space.

IV. **Figure S4:** Infiltration of microglia into the subretinal space of PRCD-deficient retina.

V. **Figure S5:** RPE atrophy in *Prcd*$^{-/-}$ mice.

VI. **Table. S1:** List of antibodies and dyes used in this study.

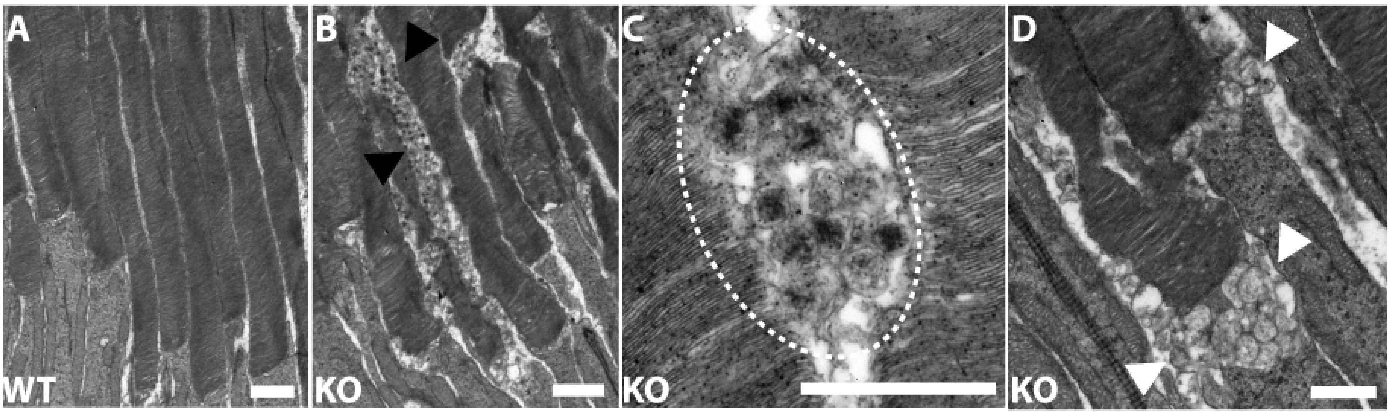


**Figure S1.** ***Prcd***$^{-/-}$ **mice accumulate EVs.** TEM images from 12-month-old (**A**) WT showing normal POS (**B-D**) *Prcd*$^{-/-}$ mice showing irregular OS disc diameters; and accumulation of EVs/ectosomes in the inter-photoreceptor matrix (**B**, *black arrowheads*; magnified in **C**, *dashed oval*) and at the base of the OS (**D**, *white arrow heads*). Scalebars – A, B: 2µm; C, D: 1µm

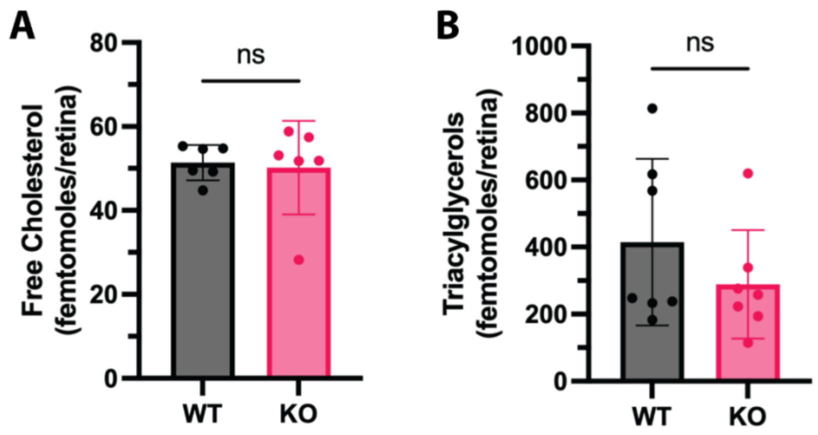


**Figure S2. FC and TAG levels are unchanged in *Prcd*$^{-/-}$ mice.** Untargeted LC-MS/MS analysis of whole retinas from *Prcd*$^{-/-}$ and WT mice at P60 showing no changes in the levels of FC (**A**) and TAGs (**B**) between the groups. Groups were compared using unpaired Student t-tests. Data are represented as mean ± SD.

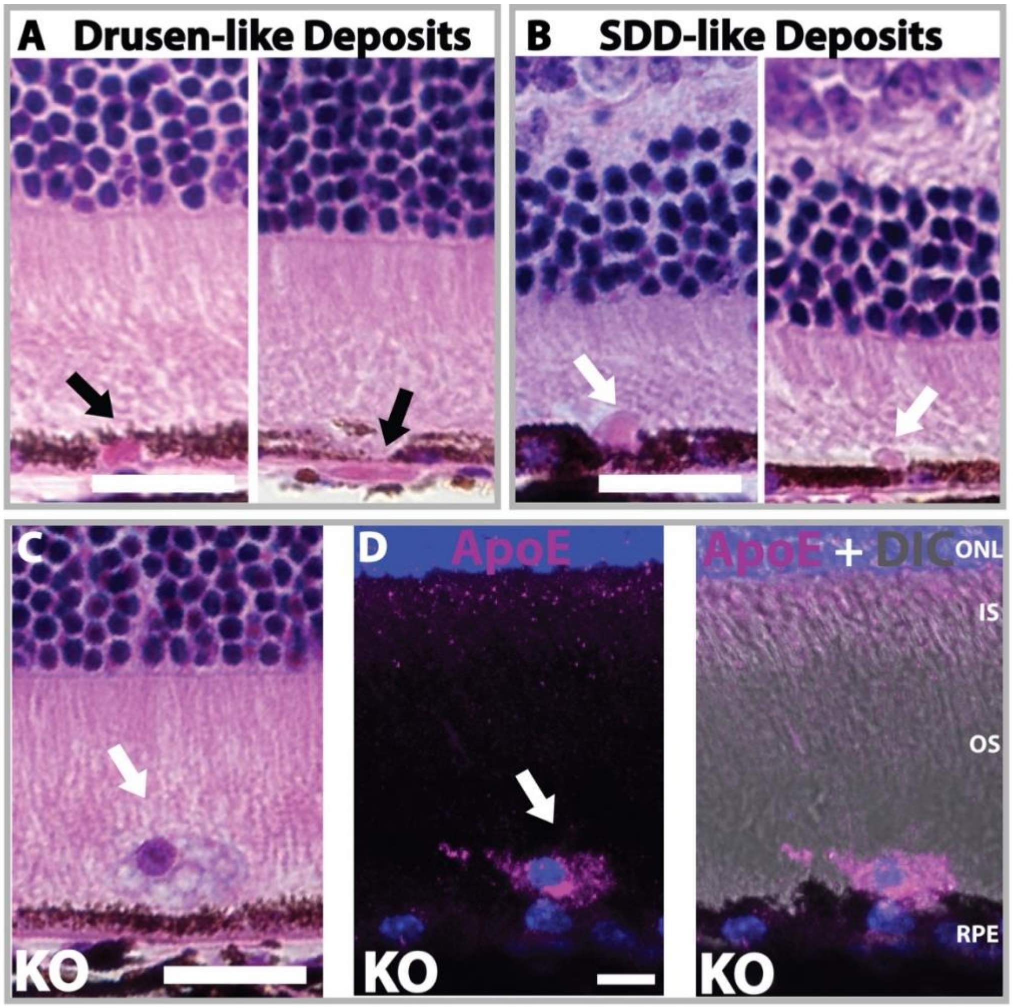


**Figure S3.** ***Prcd*$^{-/-}$ mice exhibit extracellular focal deposits and non-pigmented cells in the subretinal space.** H&E stained sections from *Prcd*$^{-/-}$ mice showing focal deposits within the RPE/BrM (**A**) and in the subretinal space (**B**). Non-pigmented cells found in the subretinal space in H&E stained sections (**C**) express ApoE (**D**). Scalebars – A,B,C: 20µm; D: 10µm

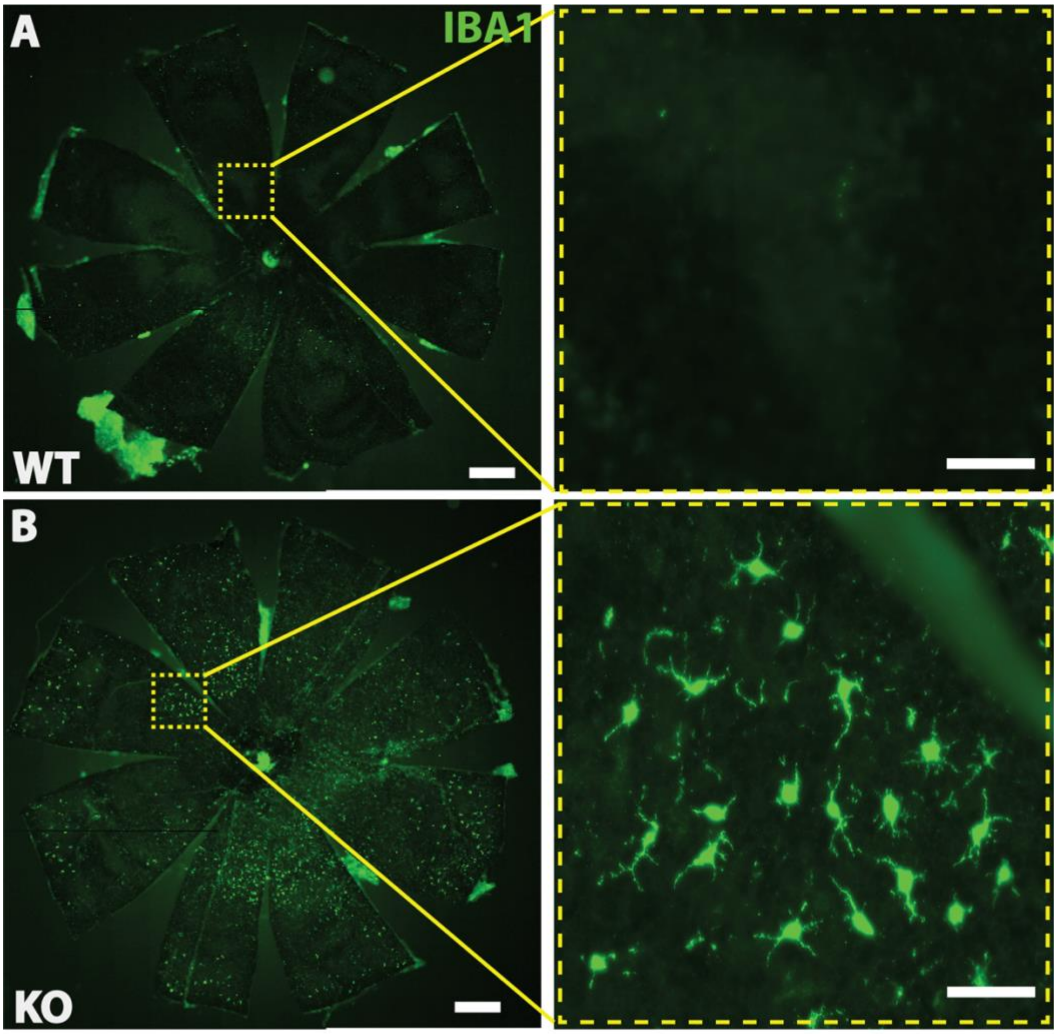


**Figure S4. Infiltration of microglia into the subretinal space of PRCD-deficient retina.** Representative images of RPE flatmounts from 12-month-old WT (**A**) and *Prcd*$^{-/-}$ mice (**B**) labelled with microglia/macrophage marker – IBA1 (*green*). Massive accumulation of subretinal microglia is observed only in *Prcd*$^{-/-}$ mice. These microglia exhibit an amoeboid phenotype suggesting an activated state (B, *magnified inset*). No microglial infiltration is observed in WT controls. Scalebars – A, B: 500µm; Magnified insets: 100µm.

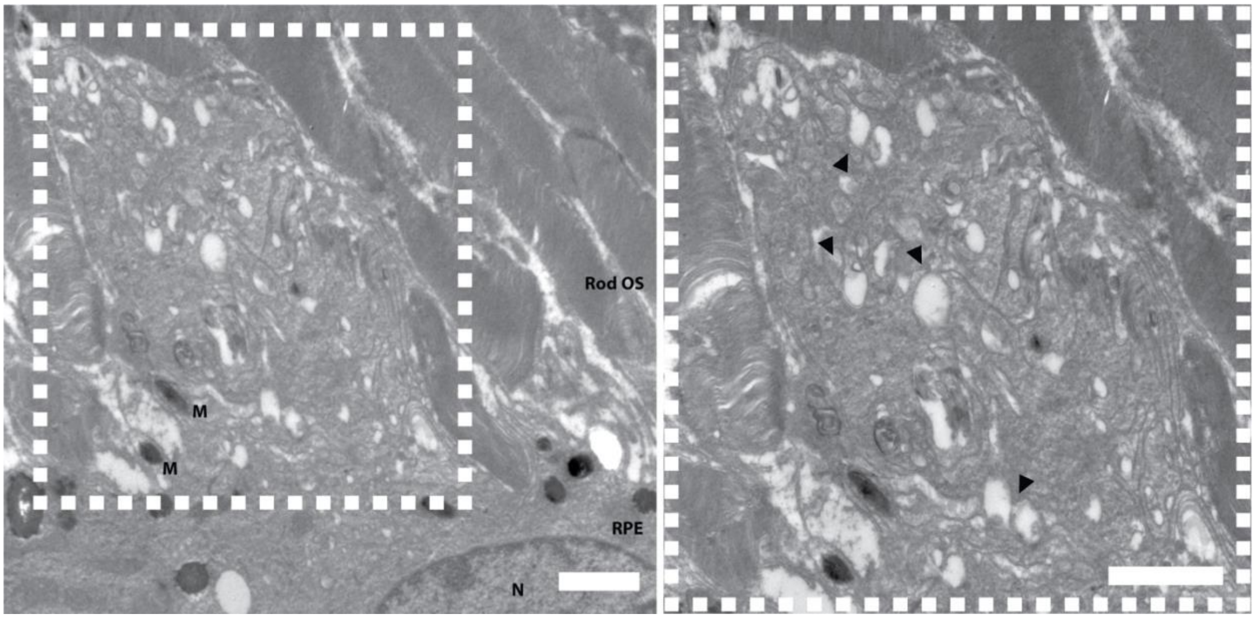


**Figure S5. RPE atrophy in *Prcd*$^{-/-}$ mice.** TEM image from 12-month-old *Prcd*$^{-/-}$ mouse showing extracellular deposits in the subretinal space resembling cellular debris from degenerating RPE. The magnified inset on the right shows accumulation of membranous vacuoles (*arrowheads*). N- Nucleus, M – Melanosome. Scalebars- 2µm.

**Table S1.** List of antibodies and dyes used in this study

| Antibody/Dye | Host | Company/ Source | Catalog # | Dilution |
|---|---|---|---|---|
| Rhodopsin | Mouse | R.Molday, Univ. British Columbia | | 1:1000 |
| Perilipin | Rabbit | Thermofisher | 15294-1-AP | 1:3000 |
| Nile Red | n/a | Sigma | N3013 | 1.75µg/mL |
| Vitronectin | Mouse | Santa Cruz | sc-74484 | 1:300 |
| ApoE | Rabbit | Protein Tech | 18254-1-AP | 1:200 |
| IBA1 | Rabbit | WAKO | 019-19741 | 1:1000 |
| Anti-mouse Alexa Fluor- 488 | Goat | Invitrogen | A11001 | 1:500 |
| Anti-mouse Alexa Fluor- 568 | Goat | Invitrogen | A11004 | 1:500 |
| Anti-mouse Alexa Fluor- 647 | Goat | Invitrogen | A32728 | 1:500 |
| Anti-rabbit Alexa Fluor 488 | Goat | Invitrogen | A32731 | 1:500 |
| Anti-rabbit Alexa Fluor 568 | Goat | Invitrogen | A11011 | 1:500 |
| 4′,6-diamidino-2-phenylindole (DAPI) | | Thermofisher | D1306 | 1:1000 |
| Alexa 488 conjugated Phalloidin | | Thermofisher | A12379 | 1:500 |
| Phalloidin-iFluor™ 647 Conjugate | | Cayman Chemical | 20555 | 1:1000 |
| Fluorescein Dye | | Altaire Pharmaceutical, Inc | NDC-59390-199-05 | 10% |
